# Spatial Transcriptomics of IPMN Reveals Divergent Indolent and Malignant Lineages

**DOI:** 10.1101/2024.10.29.620810

**Authors:** Matthew K. Iyer, Ashley Fletcher, Chanjuan Shi, Fengming Chen, Elishama Kanu, Austin M. Eckhoff, Matthew Bao, Timothy L. Frankel, Arul M. Chinnaiyan, Daniel P. Nussbaum, Peter J. Allen

**Affiliations:** Department of Surgery, Duke University; Durham, North Carolina; Department of Pathology, Duke University; Durham, North Carolina; Michigan Center for Translational Pathology, University of Michigan, Ann Arbor, Michigan; Department of Computational Medicine and Bioinformatics, University of Michigan, Ann Arbor, Michigan; Department of Pathology, University of Michigan, Ann Arbor, Michigan; Department of Surgery, University of Michigan, Ann Arbor, Michigan; Department of Urology, University of Michigan, Ann Arbor, Michigan; Howard Hughes Medical Institute, Chevy Chase, Maryland

**Author notes:** **Financial disclosures:** None. **Financial support:** R01 CA182076-07.

## Abstract

**Purpose:** Intraductal papillary mucinous neoplasms (IPMN) occur in 5-10% of the population, but only a small minority progress to pancreatic ductal adenocarcinoma (PDAC). The lack of accurate predictors of high-risk disease leads both to unnecessary operations for indolent neoplasms as well as missed diagnoses of PDAC. Digital spatial RNA profiling (DSP-RNA) provides an opportunity to define and associate transcriptomic states with cancer risk.

**Experimental Design:** Whole-transcriptome DSP-RNA profiling was performed on 10 IPMN specimens encompassing the spectrum of dysplastic changes from normal duct to cancer. Ductal epithelial regions within each tissue were annotated as normal duct (NL), low-grade dysplasia (LGD), high-grade dysplasia (HGD), or invasive carcinoma (INV). Gene expression count data was generated by Illumina sequencing and analyzed with R/Bioconductor.

**Results:** Dimension reduction analysis exposed three clusters reflecting IPMN transcriptomic states denoted “normal-like” (*cNL*), “low-risk” (*cLR*) and “high-risk” (*cHR*). In addition to specific marker genes, the three states exhibited significant enrichment for the exocrine, classical, and basal-like programs in PDAC. Specifically, exocrine function diminished in *cHR*, classical activation distinguished neoplasia from *cNL*, and basal-like genes were specifically upregulated in *cHR*. Intriguingly, markers of *cHR* were detected in NL and LGD regions from specimens with PDAC but not low-grade IPMN.

**Conclusions:** DSP-RNA of IPMN revealed low-risk (indolent) and high-risk (malignant) expression programs that correlated with the activity of exocrine and basal-like PDAC signatures, respectively, and distinguished pathologically low-grade from malignant specimens. These findings contextualize IPMN pathogenesis and have the potential to transform existing risk stratification models.

**Statement of translational relevance:** Current consensus guidelines for management of intraductal papillary mucinous neoplasms (IPMN) of the pancreas utilize clinical and radiographic criteria for risk stratification. Unfortunately, the estimated positive predictive value of these criteria for IPMN-associated pancreatic ductal adenocarcinoma (PDAC) is under 50%, indicating that over half of pancreatectomies are performed for benign disease. Moreover, nearly 15% of patients who were deemed “low risk” by the same criteria harbored PDAC. Surgical resection of IPMN has maximal benefit when performed prior to the development of PDAC, as evidence of carcinoma has been associated with a high rate of recurrence and poor overall survival. Thus, the development of molecular diagnostics that improve the accuracy of IPMN risk classification would have immediate relevance for patient care, both in terms of better selecting patients for potentially curative operations, as well as sparing patients with low-risk lesions from invasive procedures.

## INTRODUCTION

Pancreatic ductal adenocarcinoma (PDAC) portends a five-year survival rate of 12% and accounts for over 50,000 annual deaths in the United States due, in part, to the lack of effective early detection strategies^1^. Precancerous pancreatic cystic lesions known as intraductal papillary mucinous neoplasms (IPMN) account for 15-20% of PDAC cases, and are unique in that these precursors can be detected by cross-sectional imaging^2^. Current consensus guidelines utilize clinical and radiographic criteria for risk stratification of IPMN^2^. Unfortunately, the estimated positive predictive value of these criteria for malignant IPMN is under 50%, indicating that over half of pancreatectomies are performed for low-grade lesions^3^. Moreover, patients who were initially deemed “low risk” may progressed to malignant IPMN, with the rate of underdiagnosis reported at nearly 15%. Surgical resection of IPMN has maximal benefit when performed prior to the development of PDAC, as even evidence of a minute focus of carcinoma has been associated with a 24-45% risk of local or distant recurrence^4,5^. Thus, the development of molecular diagnostics that improve the accuracy of IPMN risk classification would have immediate relevance for patient care, both in terms of better selecting patients for potentially curative operations, as well as sparing patients with low-risk lesions from invasive procedures.

The discovery of tissue biomarkers to improve risk classification depends upon precise molecular profiling of IPMN specimens. Molecular characterization of normal and neoplastic pancreatic ductal epithelium has been confounded by the complex pancreatic microenvironment, including surrounding acini, islets, immune populations, and stromal components^6^. Furthermore, distinct intestinal (INT), gastric foveolar (GF), and pancreatobiliary (PB) histologic subtypes and varying grades of dysplasia often co-occur in patient specimens, limiting the specificity of bulk and single-cell profiling methods to decipher molecular differences between subtypes and grades of dysplasia^7^. Additionally, the morphologic classification and grading of IPMN tissues suffers from relatively subjective histopathologic standards, leading to inconsistencies between studies^8^. Spatial transcriptomics enables a more precise characterization of individual tissue regions, and use of these platforms by our group and others has begun to allow for translation from histopathologic to molecular subtyping^9–11^. Our previous work identified genes and biological pathways enriched in INT, GF, and PB subtypes, and showed that only PB morphology resembled PDAC. However, this work was limited by a relatively small panel of ∼1,800 genes (∼10% of the protein coding transcriptome) and did not profile normal ducts or regions of carcinoma. Subsequent work by Sans *et al.* and Agostini *et al.* defined a more comprehensive transcriptomic landscape of IPMN, but focused primarily on the molecular drivers of indolent disease^10,11^. Neither study examined the full spectrum of disease within individual patients.

In this study, we used digital spatial RNA profiling (DSP-RNA) to compare whole-transcriptomic features associated with normal ducts (NL), low-grade dysplasia (LGD), high-grade dysplasia (HGD), and invasive carcinoma (INV) within the same patient as well as between patients. As shown in lineage tracing studies of PDAC in mice, we hypothesized that transcriptomic evidence of indolent or malignant fate could be deciphered from normal or low-grade lesions^12^. We therefore aimed to define a transcriptomic classification of indolent and malignant IPMN that could be leveraged to intercept high-risk disease at an earlier stage and improve surgical decision making for patients.

## MATERIALS AND METHODS

### Specimen procurement

Patients with low-grade IPMN or IPMN-associated PDAC who underwent pancreatectomy between 3/01/2019 to 6/30/2020 were identified by reviewing the Pathology Department Archives at Duke University Health System. The Institutional Review Board (IRB) approved the use of de-identified patient specimens for retrospective molecular profiling. Informed consent was not required due to the retrospective nature of the study and de-identification of the specimens.

This study focused on patients with GF or PB lesions, building on our recent findings that intestinal-type IPMN has a unique transcriptome distinct from PDAC, whereas PB often co-occurs with GF and closely resembles PDAC^9^. Archived FFPE specimens were procured and reviewed by a board-certified pathologist specializing in pancreatic pathology to confirm diagnosis (CS). Cases were included if 1) all slides were available for pathology review, with selected block(s) available for the study, 2) they were GF or PB type and 3) both normal duct and low-grade IPMN present on the same block for the cases with low-grade IPMNs, or sufficient normal ducts, low-grade, high-grade, and invasive carcinoma present in one or two blocks for the cases with IPMN associated invasive carcinoma. Cases were excluded if 1) they were intestinal type IPMN or other variants, 2) there was equivocal high-grade dysplasia for low-grade IPMN cases, and 3) they were used by our previous study.

Ten cases were selected, including 4 low-grade IPMNs (without evidence of HGD or PDAC) and 6 with IPMN-associated PDAC. Of the cases selected, 9 were clinically diagnosed by another board-certified GI/pancreas pathologist and 1 by CS. A second board-certified GI/pancreas pathologist (FC) reviewed and confirmed the blocks and ROIs for all cases, including the case initially diagnosed by CS. Specimen blocks were cut into 5μm thick serial sections. One section was stained with hematoxylin and eosin (H&E) and imaged using a Nikon TE2000-E microscope for pathology review. A second section was mounted on a positively charged slide for spatial transcriptomics. Tissue regions for DSP-RNA were selected based on review of H&E slides.

### Digital Spatial RNA profiling

DSP-RNA profiling was conducted using the NanoString GeoMx Digital Spatial Profiler using the Whole Transcriptome Atlas (WTA) kit. Tissue slides were incubated with a cocktail of oligonucleotide probes with photo-cleavable barcodes and fluorescently conjugated antibodies: anti-pan-cytokeratin (PanCK) for epithelial cells, CD45 for immune cells, and CD3 for T cells. Regions of interest (ROIs) were drawn precisely to maximize the number of cells of identical histologic appearance while adhering to the maximum allowed size of 700um2. Segmental profiles of fluorescent antibody-stained cells within each ROI were termed Areas of Interest (AOIs). Libraries were prepared according to the NanoString GeoMx Library Preparation Manual and pooled to equimolar concentration. RNA was sequenced under standard conditions on an Illumina Novaseq 6000 to a depth of 30 read-pairs/μm2. The GeoMx DSP Analysis Server performed trimming, alignment, and deduplication of barcodes using their unique molecular identifiers (UMIs). A tabulated matrix of UMI probe counts for each AOI was exported from the DSP server for subsequent analysis.

### Quality control and normalization

Quality control (QC) metrics for each AOI were computed as previously described^9^. Briefly, a receiver operating characteristic (ROC) analysis was used to obtain the signal-to-noise area under the curve (snAUC) as a measure of AOI quality, and a limit of detection (LOD) for each AOI was computed as the 90^th^ percentile of the background probe counts^13^. The following quality control criteria were used to filter AOIs: 1) >30,000 unique molecular identifier (UMI) counts, 2) snAUC > 0.60, and 3) >20% of gene probes above the LOD. AOIs that did not meet these QC criteria were removed. Background probe levels were used to estimate the false positive rate for declaring a gene expressed as well as area under the curve (AUC) for gene versus background probes. Gene probes above the LOD in at least >15% of AOIs were considered expressed. Background correction was performed to minimize noise variation across AOIs. Background-subtracted counts were scaled by UMI counts and subjected to quantile normalization^14^.

### External Validation Datasets

Nanostring GeoMx DSP data from Agostini *et al.* (GSE229752) and Carpenter *et al*. (GSE226829) were downloaded from the Gene Expression Omnibus (GEO)^11,15^. The Nanostring GeoMx Spatial Organ Atlas dataset was obtained from the company website (https://nanostring.com/resources/hu_pancreas_workflow_and_count_files-tar-gz/). The unprocessed probe count data were subjected to the same quality control and normalization procedures described above.

### Dimension reduction analysis and clustering

Analysis of filtered, normalized gene expression data was performed in R/Bioconductor^16^. Highly variable genes were determined by ranking the ratio between the observed and expected variance determined by fitting a local polynomial regression (LOESS) curve to the mean-variance relationship. Principal component analysis (PCA) utilized the 500 most highly variable genes in the entire dataset. Unsupervised clustering of principal components was performed with Partitioning Around Medioids (PAM) using the Euclidean distance metric. The average silhouette method was used to select the optimal number of clusters.

### Differential expression analysis

Differential expression (DE) analysis was conducted using the *limma* package^17^. Initial criteria for calling DE genes included absolute log2 fold change > 1.0 and adjusted p-value < 0.05. Stricter criteria (absolute log2 fold change > 3.0, adjusted p-value < 0.01) were enforced when calling marker genes. Heatmap plots were generated using the *pheatmap* R package (https://cran.r-project.org/web/packages/pheatmap/index.html). Hierarchical clustering of heatmap rows and columns used the “ward.D2” method.

### External Gene Sets

RNA-Seq analysis results from the National Cancer Institute’s Clinical Proteomic Tumor Analysis Consortium (CPTAC) proteogenomic characterization of pancreatic adenocarcinoma were obtained from Supplementary Table S3 of the published manuscript^18^. Genes were merged by official gene symbol. Differentially expressed RNAs and proteins between PDAC and benign adjacent tissue with an absolute log2 fold change > 1.0 and adjusted p-value < 0.05 were included in gene sets. Human Protein Atlas (HPA) pathology data were downloaded from the HPA web portal (https://www.proteinatlas.org/download/pathology.tsv.zip) and joined by gene symbol^19^. Pancreatic cancer molecular subtype gene sets were downloaded from the open source *pdacR* package (https://github.com/rmoffitt/pdacR) and joined by gene symbol with the filtered normalized data^20^. Hallmark gene sets were obtained from the Molecular Signatures Database (MSigDB) version 2023.2^21^. A table of the gene sets utilized in this study is provided (**Supplementary Table S1**).

### Gene Set Enrichment Analysis

Gene Set Enrichment Analysis (GSEA) was conducted using the Bioconductor package *fgsea*^22^. Ranked gene lists were weighted by log2 fold change from the associated DE comparison. For each gene set, the most significant result (smallest adjusted p-value) between positive and negative one-tailed enrichment scores was used. Adjusted p-values < 0.01 were considered significant. Leading edge genes were extracted from GSEA and DE result tables and plotted using the *pheatmap* package. Gene Set Variation Analysis (GSVA) was used with default parameters to score individual AOIs^23^.

### Trajectory inference

Trajectory inference was performed using the Bioconductor package *slingshot* with default parameters^24^. Inputs included the matrix of PCA results, a vector of AOI cluster labels, and designated start (*cNL*) and end (*cHR*) clusters for lineage mapping.

## RESULTS

### Spatial transcriptomic profiling of low-grade and malignant IPMN

We performed DSP-RNA to examine transcriptomic states associated with IPMN progression in a cohort of ten patients, including six with IPMN-associated cancer (PDAC specimens) and four with only low-grade dysplasia (LG specimens) (**Table 1**, **Supplementary Table S2, Fig. 1A**). A pancreatic pathologist guided the selection of regions of normal duct (NL), low-grade dysplasia (LGD), high-grade dysplasia (HGD), and invasive carcinoma (INV) from each specimen. Examples from a patient with IPMN-associated cancer (*p10*) illustrate the study design (**Fig. 1B**, **Supplementary Fig. S1**). Fluorescent antibody segmentation was utilized to profile 104 epithelial (PanCK+), 42 immune (CD3+, CD45+), and 42 stromal (PanCK-, CD3-, CD45-) AOIs. The median nuclei count per AOI was 351 (IQR 116-633). High-throughput sequencing yielded a median of 284,042 UMI counts per AOI (IQR 62,571 – 732,588). Total UMI counts were correlated with AOI nuclei count (Pearson r=0.70, p < 2.2e-16) and surface area (Pearson r=0.45, p=6.2e-11), supporting an association with *in situ* RNA abundance (**Supplementary Fig. S2A, B**)^9^. Epithelial and stromal AOIs, which exhibited greater cellularity, yielded higher quality transcriptomic profiles than immune AOIs (**Supplementary Fig. S2C, D, E**). Within the epithelial segment, neoplastic AOIs (LGD, HGD, INV) produced richer profiles than normal ducts (**Supplementary Fig. S2F, G, H)**.

**Figure 1:**
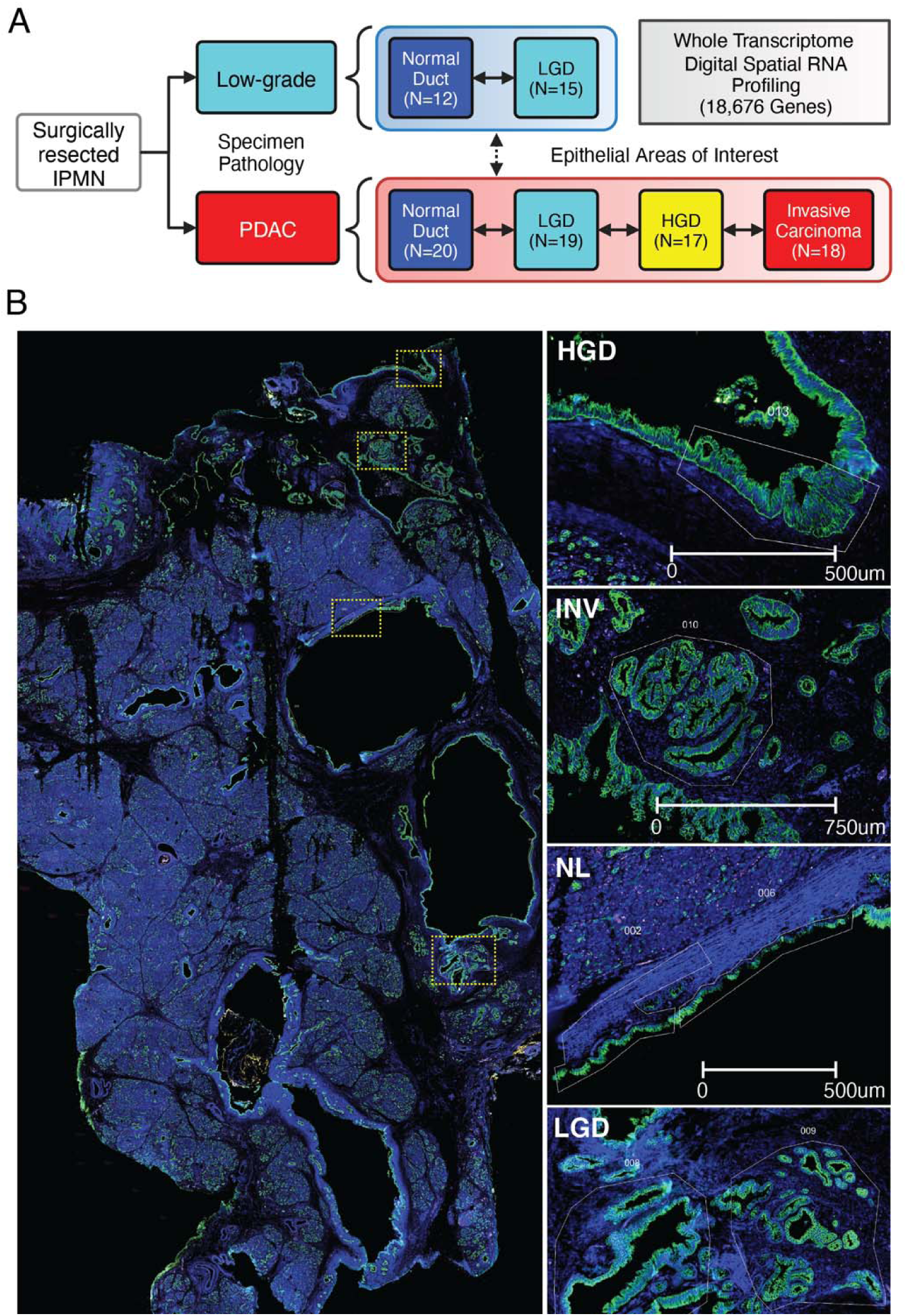
Digital Spatial Profiling of PB-IPMN. **A)** Study design diagram, **B)** Tissue slide from patient *p10* stained for PanCK (green), CD45 (yellow), CD3 (red), and DNA (blue). Highlighted boxes showing examples of normal (NL), low-grade (LGD), high-grade (HGD), and invasive carcinoma (INV) areas of interest (AOIs) selected for digital spatial profiling.

**Table 1:** Patient characteristics of IPMN spatial transcriptomics cohort.

|  |  |
| --- | --- |
| Patients | 10 |
| Median age: years (IQR) | 74.5 (range 52-77) |
| Female Sex | 5 (50%) |
| Race |  |
| Caucasian | 9 (90%) |
| Asian | 1 (10%) |
| Black | 0 (0%) |
| Unidentified | 0 (0%) |
| Operation |  |
| Pancreaticoduodenectomy | 7 (70%) |
| Distal pancreatectomy | 3 (30%) |
| Other | 0 (0%) |
| Median lesion size | 3.9cm (range 2.4-5.5cm) |
| Invasive carcinoma within lesion | 6 (60%) |
| Invasive component size | 2.65cm (range 2.0-5.5cm) |
| Cancer stage | IB (2), IIA (2), IIB (2) |
| Cancer recurrence | 3 (50%) |
| Death | 3 (30%) |
| Anatomic subtype |  |
| Main duct | 2 (20%) |
| Branch duct | 3 (30%) |
| Mixed | 2 (20%) |
| Indeterminate | 3 (30%) |
| Dominant histologic subtype |  |
| Intestinal | 0 |
| Pancreatobiliary | 6 |
| Gastric foveolar | 4 |

Filtering based on QC criteria retained 101 of 104 epithelial (97%), 34/42 stromal (80%), and 11/42 immune (26%) AOIs (**Supplementary Fig. S3A-C**). Due to their poor technical quality, immune AOIs were omitted from subsequent analysis. Epithelial and stromal data subsets were analyzed separately. Gene probes indistinguishable from background noise were removed prior to data normalization, resulting in 12,451 (67%) and 10,399 (56%) expressed genes in the epithelial and stromal subsets, respectively (**Supplementary Fig. S4**). Our normalization approach effectively mitigated the effect of background noise on gene expression variation (Methods, **Supplementary Fig. S5**).

### Three major states comprise the epithelial transcriptome of IPMN

To discover epithelial states that comprise IPMN in an unsupervised manner, we employed principal component analysis (PCA). The first two principal components (PCs) explained 34.77% of the variance in the dataset, with a substantial decline (15.37% to 6.71%) in explained variance occurring from PC3 onward (**Supplementary Fig. S6A**). Technical variation in background noise did not contribute to the top PCs, suggesting that the results reflected biological sources of gene expression variation (**Supplementary Fig. S6B**). Indeed, visualization of the top genes contributing to PC1 and PC2 depicted known markers of IPMN (*MUC5AC*) and PDAC (*TRIM29*) (**Supplementary Fig. S6C**)^25,26^. Accordingly, the first two PCs effectively delineated distinct groups of LGD and NL AOIs that diverged from the HGD and INV AOIs (**Fig. 2A,B**). In addition, the PCA plot highlighted specimens with prominent intratumoral heterogeneity (**Fig. 2C**). For example, AOIs from patient *p04* were distributed widely across the PC1 and PC2 axes. In addition, *all* the AOIs from patient *p07* were skewed towards the positive end of PC1, except for a single LGD AOI.

**Figure 2:**
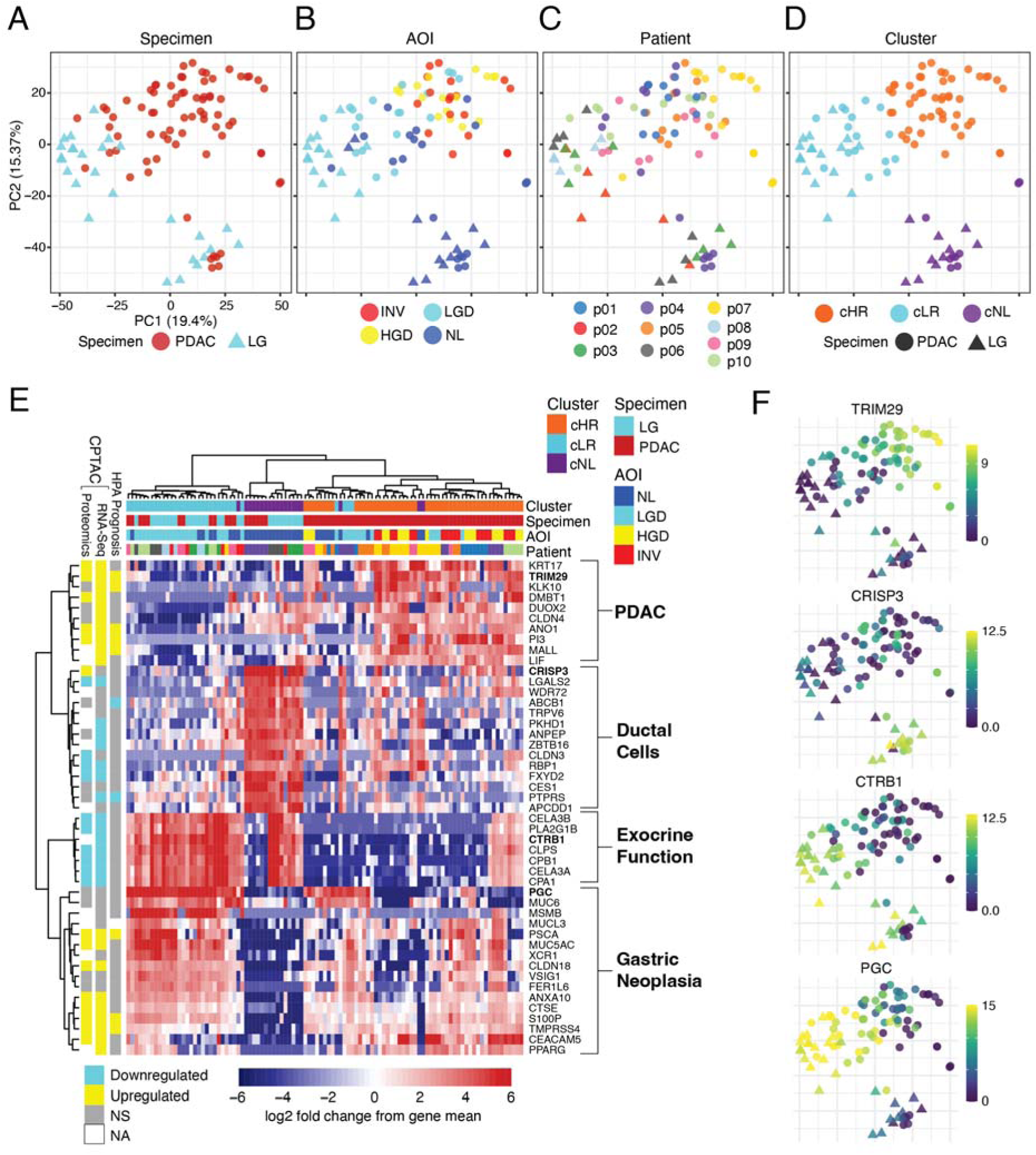
Unsupervised analysis to define transcriptomic states of IPMN. **(A-D)** Scatter plot of the first two Principal Components obtained from filtered, normalized spatial transcriptomic data colored by **(A)** specimen pathology, **(B)** AOI pathology, **(C)** patient, and **(D)** cluster. **(E)** Heatmap plot of marker genes obtained from pairwise comparison across clusters (*cHR-vs-cLR*, *cHR-vs-cNL*, *cLR-vs-cNL*) with hierarchical clustering of rows (AOIs) and columns (genes). Gene expression is shown as log2 fold change from the gene median across all samples. Genes are annotated by evidence of upregulation/unfavorable prognosis or downregulation/favorable prognosis in PDAC according to Clinical Proteomic Tumor Atlas Consortium (CPTAC) or Human Protein Atlas (HPA) data. **(F)** PCA scatter plots colored by the expression of select marker genes *TRIM29*, *CRISP3, CTRB1*, and *PGC* across all AOIs.

To categorize the gene expression patterns revealed by PCA, we employed unsupervised clustering to define three transcriptomic states, after confirming that a three-cluster model was optimal for the dataset (**Fig. 2D**, **Supplementary Fig. S6D**). The three clusters exhibited strong enrichment for pathologic grade (X^2^ (df=6, N=101) = 102.85, p < 2.2e-16, **Supplementary Table S3**). One cluster was composed of *only* NL AOIs and was therefore designated “normal-like” (*cNL*). A second cluster consisted of *only* NL and LGD AOIs and was labeled “low-risk” (*cLR*). The remaining cluster harbored a mix of pathologic grades, including HGD and INV AOIs, and was labeled “high-risk” (*cHR*). Intriguingly, *all* AOIs within *cHR* were from patients with PDAC.

### Identification of marker genes associated with transcriptomic states

Pairwise DE analysis of the AOIs within *cNL*, *cLR*, and *cHR* clusters was performed (*cLR-vs- cNL*, *cHR-vs-cNL, cHR-vs-cLR*) to identify marker genes associated with transcriptomic states (**Supplementary Table S4)**. For each comparison, the top genes that met strict thresholds were nominated as marker genes (top 10 overexpressed/underexpressed genes, absolute log2 fold change > 3, adjusted p-value < 0.01). The union of these comparisons resulted in 47 marker genes (**Fig. 2E**). Marker genes were annotated by their differential expression status in the CPTAC bulk RNA-Seq analysis of PDAC versus benign adjacent tissue and by their association with PDAC prognosis in the Human Protein Atlas (HPA)^18,19^. Examination of the hierarchical clustering of marker genes, their annotations, and literature evidence revealed four groups with clear biological relevance: 1) pancreatic exocrine function (ex. *CTRB1*), 2) gastric neoplasia (ex. *PGC, MUC6*), 3) normal pancreatic ducts (ex. *CRISP3*), and 4) PDAC (ex. *KRT17, TRIM29*)^10,27^. Marker genes within the pancreatic exocrine (7/7, 100%) and ductal epithelial (8/14, 57%) groups were often downregulated in PDAC, whereas marker genes within the gastric neoplastic (9/16, 56%) and PDAC (10/10, 100%) groups were upregulated in PDAC. Plotting the expression of exemplary genes from each group on the PCA axes delineated the three transcriptomic states (**Fig. 2F**).

### Transcriptomic states of IPMN associate with PDAC molecular subtypes

To contextualize IPMN spatial expression patterns, we compared our DE results to published PDAC bulk RNA-Seq gene sets using GSEA (**Fig. 3A**, **Supplementary Table S5**). The enrichment scores for upregulated and unfavorable prognosis genes in PDAC showed a stepwise increase across the *cLR-vs-cNL*, *cHR-vs-cNL*, and *cHR-vs-cLR* contrasts, respectively. For downregulated or favorably prognostic genes in PDAC, *cHR-vs-cNL* and *cHR-vs-cLR* produced greater negative scores than *cLR-vs-cNL*. Collectively, the GSEA results supported our IPMN classification scheme, particularly the association of *cHR* with malignancy.

**Figure 3:**
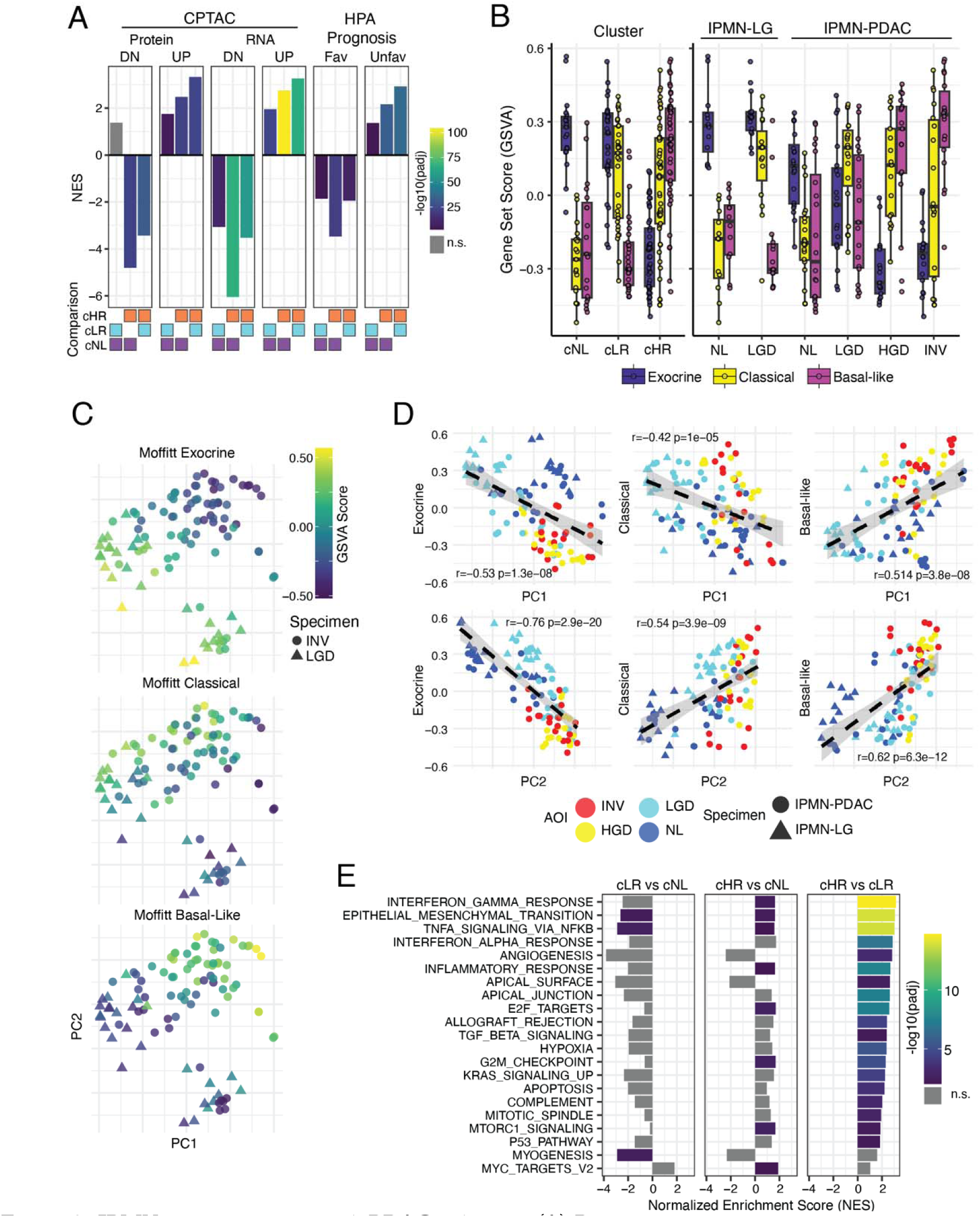
IPMN states associate with PDAC subtypes. **(A)** Barplots showing the normalized enrichment score (NES) from gene set enrichment analysis (GSEA) of *cHR-vs-cLR*, *cHR-vs-cNL*, and *cLR-vs-cNL* comparisons across PDAC gene sets from CPTAC and HPA. Bars are colored by the adjusted p-value obtained from GSEA, or grey if not significant. **(B)** Boxplots comparing gene set variation analysis (GSVA) for Moffitt Exocrine, Classical, and Basal-like gene signatures. **(C)** PCA scatter plots colored by GSVA scores for the Moffitt Exocrine, Classical, and Basal-like subtypes. **(D)** Scatter plots comparing Principal Components (*x* axis) to Moffitt gene set scores (*y* axis) for PC1 and PC2. Plots are annotated with Pearson correlation coefficients and p-values. **(E)** Barplots of GSEA results for significant MSigDB Hallmark Pathways. Bar colors reflect degree of significance. DN=Down, Fav=Favorable Prognosis, Unfav=Unfavorable Prognosis.

Considering the modest yet significant enrichment for PDAC gene sets in *cLR*, we hypothesized that some PDAC expression programs could be common to *cLR* and *cHR*, while others might be specific to each state. To investigate the possibility, we tested whether our IPMN clusters were enriched for the established molecular subtypes of PDAC defined by Moffit *et al*., Collison *et al.*, Bailey *et al*., and Chan-Seng-Yue *et al.* (**Supplemental Fig. S7**)^20,28–31^. Three clear patterns were evident from this GSEA analysis. First, the classical subtype of PDAC (Moffitt Classical, Collison Classical, ICGC Immunogenic Up, Chan Seng Yue Classical A) was positively enriched in both *cHR-vs-cNL* and *cLR-vs-cNL*. This suggests that both *cLR* and *cHR* upregulate classical PDAC genes relative to *cNL*. Interestingly, testing *cHR-vs-cLR* produced significant negative scores (Moffitt Classical, ICGC Immunogenic Up, and Chan-Seng-Yue Classical A), suggesting that *cLR* upregulated classical pathway genes to a greater degree than *cHR*. Second, pancreatic exocrine function (Moffitt Exocrine, Collison Exocrine, and ICGC ADEX) was markedly diminished in *cHR-vs-cNL* and *cHR-vs-cLR*, suggesting that *cHR* epithelium lacks exocrine function, whereas *cLR* retains some exocrine function. Third, positive enrichment of basal-like genes (Moffit Basal-like, Collison Quasi-mesenchymal, Chan-Seng-Yue Basal A and B, ICGC Squamous) occurred exclusively in *cHR-vs-cNL* and *cHR-vs-cLR*. By contrast, *cLR-vs- cNL* produced negative or non-significant enrichment scores for basal-like signatures. The concurrent activation of basal-like genes and loss of exocrine expression was a powerful discriminator of the *cHR* state.

The integration of multiple independent studies of PDAC highlighted exocrine, classical, and basal-like genes that distinguished IPMN subtypes. However, the differences in gene set size and composition among molecular subtyping studies led us to adopt the Moffitt *et al.* gene sets for further analyses of PDAC subtypes for several reasons: 1) our DE results were most highly enriched for the Moffitt subtypes by GSEA, 2) subsequent work by Moffitt *et al.* exposed issues related to tumor purity and stromal contamination within the ICGC and Collisson gene sets, and 3) our dataset did not demonstrate the presence of Classical A/B and Basal-like A/B subtypes as in Chan-Seng-Yue *et al.*^20^ We then computed Moffitt Exocrine, Classical, and Basal-like scores for individual AOIs using Gene Set Variation Analysis (GSVA) and examined patterns of PDAC subtype expression across clusters and pathologic annotation (**Fig. 3B**)^23^. Across clusters, exocrine activity was high in *cNL* and *cLR* and diminished in *cHR*. By pathology, exocrine gene expression was higher in LG than PDAC specimens and relatively low in HGD and INV AOIs. Classical gene expression was upregulated in both *cLR* and *cHR* relative to *cNL*. Interestingly, the variation in classical scores was high within INV AOIs, alluding to the potential for classical/basal-like phenotype scoring to stratify patients with PDAC^32^. By contrast, basal-like expression was highest in *cHR* and further stratified HGD and INV AOIs. Clear exocrine, classical, and basal-like expression patterns could be visualized along the first two PCs (**Fig. 3C**). While exocrine scores exhibited a negative correlation with PC1 and PC2, basal-like activity demonstrated a positive correlation with these PCs (**Fig. 3D**). Thus, an exocrine-basal-like axis accounted for the largest portion of gene expression variation in the entire dataset. Classical expression, which was negatively correlated with PC1 and positively correlated with PC2, differentiated “normal-like” (*cNL*) from neoplastic (*cLR* and *cHR*) epithelium.

To highlight individual genes contributing to the significant enrichment of PDAC subtypes, we performed leading-edge analysis of the exocrine, classical, and basal-like gene signatures^33^. Genes that were DE and enriched within the leading edge at least two PDAC molecular subtyping studies were included in consensus gene subsets. We also associated the leading-edge genes with published PDAC bulk RNA-Seq gene sets as an additional means of validation. The exocrine leading edge included 50 genes, of which 41 (82%) were downregulated at the RNA or protein level and none were prognostic in PDAC (**Supplementary Fig. S8A**). The classical leading edge included 83 genes, of which 48 (57%) were upregulated in PDAC but only 4 (5%) were associated with unfavorable prognosis (**Supplementary Fig. S8B**). Of note, 52/83 classical subtype genes were negatively enriched in *cHR-vs-cLR*, supporting the observation that the classical program may be activated to a greater extent in *cLR*. Among the negatively enriched genes in *cHR-vs-cLR* was *GATA6*, which has been previously shown to distinguish classical from basal-like PDAC^34^. Finally, the basal-like leading edge featured 81 genes, of which 45 (55%) were supported by PDAC bulk RNA-Seq or proteomics, and 36 (44%) were associated with unfavorable prognosis (**Supplementary Fig. S8C**).

### Hallmark molecular pathways distinguish high-risk from other regions of IPMN

We assessed molecular pathway activity in IPMN by performing GSEA using the Hallmark gene set collection from the Molecular Signatures Database (MSigDB)^21^. The most enriched pathways in *cHR-vs-cLR* included interferon gamma (IFN-γ), epithelial to mesenchymal transition (EMT), and tumor necrosis factor signaling via nuclear factor κB (TNF-NFκB) (**Fig. 3E**). The involvement of inflammatory signaling in progression of IPMN was highlighted by prior work from our group and others^9,11,35^. Consistent with the strong enrichment seen in *cHR-vs-cLR*, IFN-γ, TNF-NFκB, and EMT pathways were positively enriched in *cHR-vs-cNL* and either negatively enriched or not-significant in *cLR-vs-cNL*. Within clusters and pathologic annotations, the activity of TNF-NFκB, IFN-γ, and EMT exhibited considerable variability (**Fig. S9A**). The most consistent observation was quiescent activity of IFN-γ, TNF-NFκB, and EMT activity in *cLR*. In fact, LGD AOIs from low-grade IPMN demonstrated even lower pathway activity levels than their corresponding NL counterparts.

To explore the relationship between inflammatory signaling, IPMN, and basal-like PDAC, we performed leading edge analysis of the enriched genes from the IFN-γ, TNF-NFκB, and EMT pathways (**Fig. S9B**). The basal-like genes *AREG*, *LAMC2, LAMA3, ITGA2,* and *COL7A1* were enriched within the EMT pathway. Intriguingly, *AREG* was one of three genes (along with *LAMB3* and *DUSP5*) associated with both EMT and TNF-NFκB signaling. One of the epithelial growth factor receptor (EGFR) ligands, *AREG* has been widely studied in multiple human cancers, including pancreatic cancer^36^. Additionally, *LAMC2* has been implicated in EMT and studied as a potential therapeutic target as well as a blood-based biomarker for early detection of PDAC^37,38^. Focusing on these and other enriched genes in basal-like PDAC may guide investigation towards specific therapeutic vulnerabilities.

### Stromal signatures associate with low-risk and high-risk epithelial states

For a subset of epithelial AOIs we generated DSP-RNA profiles of the adjacent stroma to explore patterns of epithelial-stromal coexpression. This yielded 34 high-quality AOIs after QC filtering, including 30 from IPMN-PDAC and 4 from IPMN-LG due to the sparsity of stromal proliferation in low-grade specimens. Gene expression variation was dominated by PC1 (20.6%), which explained nearly as much variance as PC2 and PC3 combined (21.9%) (**Supplementary Fig. S10A)**. Genes contributing to PC1 and PC2 included acinar/exocrine cell markers (*CTRB1*), islet cell markers (*INS*), neoplastic markers (*PGC, MUC6*) and published markers of activated stroma in PDAC (*COL10A1*, *POSTN*)^28^. This indicated that our stromal AOIs contained a mixture of cell types which was expected due to the lack of specific staining and segmentation of these regions (**Supplementary Fig. S10B)**^28^. To isolate stromal expression patterns, we performed PCA using highly variable genes that also occurred in the Moffitt “Normal Stroma” and Moffitt “Activated Stroma” signatures (76 genes, **Supplementary Table S6**, **Supplementary Fig. S10C**). The value of PC1 was significantly associated with transcriptomic state (χ^2^ (df=2, N=31) = 9.76, p=0.0076), but not the histologic grade (χ^2^ (df=2, N=31) = 4.40, p=0.11) of the adjacent epithelium. We pursued this compelling result further by testing for gene set enrichment after stratifying stromal AOIs by their corresponding epithelial transcriptomic states. Relative to *cLR*, the *cHR* state was negatively enriched for “Normal Stroma” (NES=-2.13, padj=2.93e-05) and positively enriched for “Activated Stroma (NES=4.41, padj=4.32e-20) (**Supplementary Fig. S10D**). Conversely, *cLR* was significantly depleted of “Activated Stroma” gene expression (NES=-4.20, padj=3.56e-14). Scoring individual AOIs using GSVA confirmed the pattern of “Normal Stroma” downregulation and “Activated Stroma” upregulation in malignant AOIs when stratified by transcriptomic state or histologic grade (**Supplementary Fig. S10E**). By contrast, the “Activated Stroma” signature was notably downregulated in LGD AOIs.

### Transcriptomic signatures stratify normal ducts and low-grade dysplasia into low-risk and high-risk groups

Although NL and LGD AOIs from low-grade and malignant specimens were histologically indistinguishable, we observed clear differences in their gene expression patterns. This suggested that the adjacent “field cancerization” associated with malignant foci could be incorporated into a gene expression biomarker^39^. To explore this possibility in an unbiased manner, we removed HGD and INV AOIs from the dataset and reanalyzed *only* NL and LGD AOIs from each patient. In this subset, PCA successfully separated AOIs from low-grade and malignant specimens and delineated a group of NL AOIs (**Fig. 4A-C**). Unsupervised clustering of NL-LGD AOIs mirrored the three-cluster assignments of the full dataset, except for seven *cNL* or *cLR* AOIs that were reclassified as *cHR* (**Fig. 4D, E)**. A 2D scatter plot representation of the pairwise DE landscape (*cHR-vs-cNL*, *cLR-vs-cNL*, and *cHR-vs-cLR*) highlighted similarities and differences between the clusters (**Supplementary Table S7**, **Fig. 4F,G**). Global DE patterns between *cHR-vs-cNL* and *cLR-vs-cNL* were highly correlated (Pearson correlation coefficient = 0.71, p-value < 2.2e-16). Indeed, loss of ductal markers (*CRISP3*, *CLDN3*, and others) and gain of neoplastic markers (*S100P, CLDN18*, and others) was common to *cLR-vs-cNL* and *cHR-vs-cNL*. However, significant differences between *cLR* and *cHR* were also evident. First, gastric neoplasia markers (*PGC*, *MUC5AC*) were downregulated in *cHR* relative to *cLR*. Second, pancreatic exocrine function (*CTRB1*, *CPA1*, *CPB1*) was lost in *cHR* AOIs. Third, certain PDAC-associated markers (*CLDN4*, *DUOX2*) were preserved in *cHR* but lost in *cLR*. Fourth, a group of “high-risk” marker genes were specifically upregulated in *cHR* relative to *cLR* and *cNL* AOIs, including *TNS4, NNMT*, and others. Together, a set of 25 top marker genes from the pairwise DE analysis of NL- LGD ROIs supported three distinguishing features of the *cHR* group: 1) loss of ductal cell markers, 2) loss/absence of exocrine function, and 3) maintenance or upregulation of specific PDAC-associated markers (**Fig. 4H**). GSEA of the NL-LGD pairwise DE results faithfully recapitulated the enrichment patterns from the full dataset for PDAC-associated genes, molecular subtypes, and Hallmark molecular pathways (**Supplementary Fig. 11A, B, C**, **Supplementary Table S8**).

**Figure 4:**
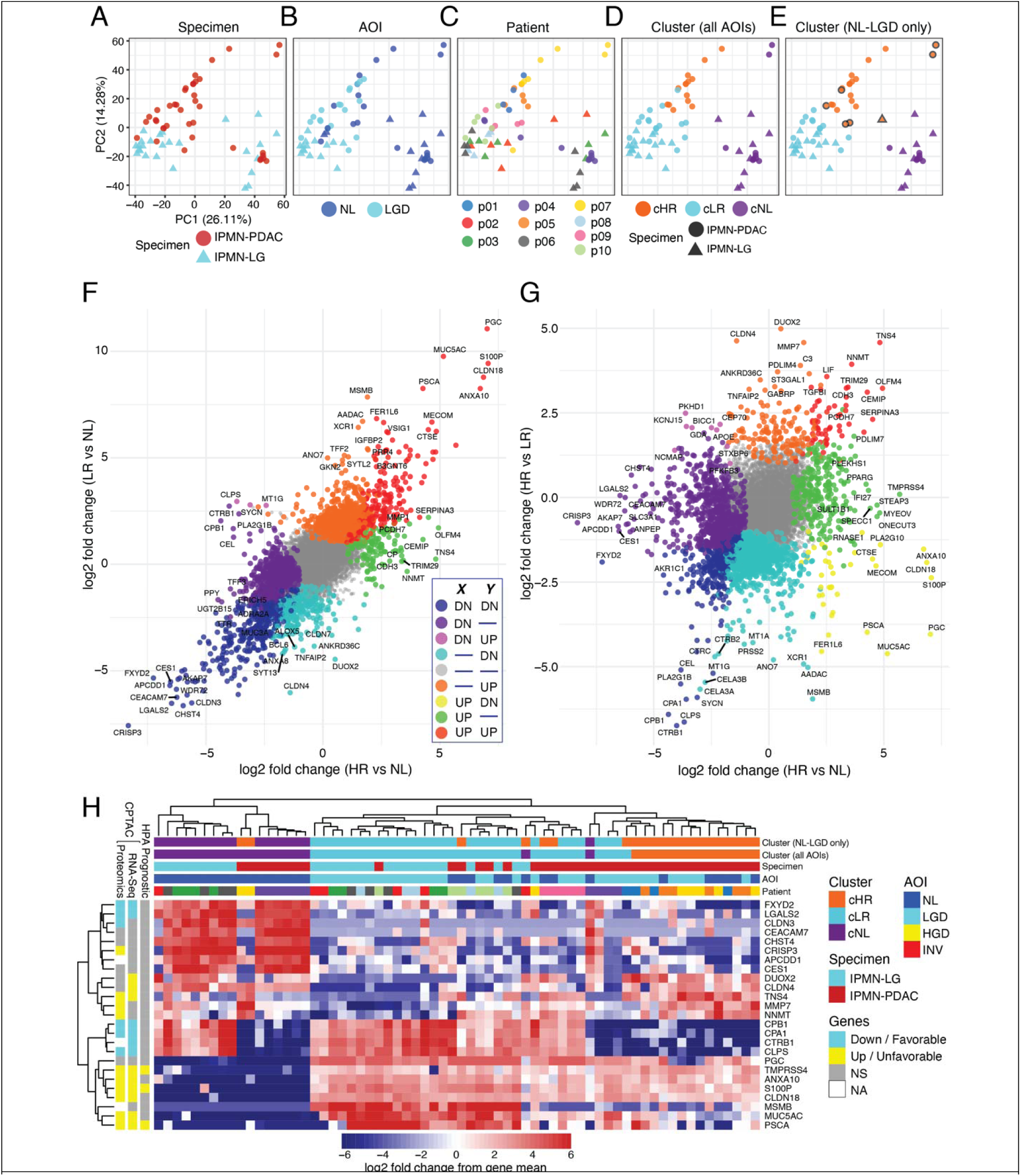
Malignant progenitor signature detectable in normal ducts and low-grade dysplasia. **(A-E)** PCA of NL and LGD AOIs from low-grade and malignant specimens colored by **(A)** specimen pathology, **(B)** AOI pathology, **(C)** patient, **(D)** cluster assignment from full dataset, and **(E)** cluster assignment from analysis of *solely* NL and LGD AOIs. Points with black outlines represent AOIs reclassified as *cHR* **(F, G)** scatter plot of two DE comparisons with log2 fold change shown on both *x* and *y* axes. Colors assigned to significantly DE genes based on upregulation/downregulation in one or both comparisons. Grey points are not significantly DE in either analysis. **(F)** *cHR-vs-cNL* (*x* axis) versus *cLR-vs-cNL* (*y* axis). **(G)** *cHR-vs-cNL* (*x* axis) versus *cHR-vs-cLR* (*y* axis). **(H)** heatmap plot showing the top 5 upregulated and downregulated marker genes from each comparison of NL-LGD clusters.

### Independent validation datasets support a generalizable three-state transcriptomic model for pancreatic cancer precursors

To assess the generalizability of the three-state transcriptomic model for IPMN, we obtained and processed three external Nanostring GeoMx WTA datasets from 1) the publicly available Spatial Organ Atlas (SOA) which profiled 24 areas of normal pancreatic ductal epithelium (PanCK+) from 4 patients, 2) the Agostini *et al.* Tissue Micro Array study of 80 epithelial (PanCK+) AOIs from 40 patients with IPMN (GSE229752), and 3) the Carpenter *et al*. study of early neoplastic lesions in deceased donor pancreas and PDAC specimens (GSE226829)^11,15^. Our processing and QC pipeline retained 24/24 (100%) of the SOA AOIs, 178/197 (90%) of the Carpenter AOIs, but just 21/80 (26.25%) of the Agostini AOIs, largely due to excessive background noise (**Supplementary Fig. S12**). Of note, the Agostini *et al*. study also used a three-tier dysplasia classification that included LGD (N=1), intermediate grade/borderline dysplasia (IGD, N=5), and HGD (N=15)^8,40^.

We then projected each of the validation AOIs (test data) onto the principal component axes defined by our IPMN AOIs (training data) and assigned them to clusters (**Supplementary Table S9**). All normal ductal AOIs (24 out of 24) from SOA clustered within *cNL*, confirming that *cNL* indeed represents normal epithelium (**Supplementary Fig. S13A, B**). By contrast, only one of the Agostini IPMN AOIs grouped with *cNL*, and the remaining 20 out of 21 Agostini AOIs were assigned to neoplastic states but did not clearly distinguish *cLR* from *cHR*.

Examination of the PCA projections of normal ducts, pancreatic intraepithelial neoplasia (PanIN), and PDAC from the Carpenter *et al*. dataset provided compelling support for our three-state transcriptomic model of pancreatic cancer precursors (**Supplementary Fig. S13C, D**). Normal ducts, PanIN, and PDAC were significantly enriched within *cNL*, *cLR*, and *cHR*, respectively (χ^2^ (df=4, N=178) = 194.4, p < 2.2e-16, **Supplementary Table S9**). Importantly, *none* of the normal ducts or incidental PanINs from donor pancreata were assigned to *cHR*, reinforcing that *cHR* represents a malignant lineage. However, some normal ducts (12/28, 42%) and PanINs (4/53, 7.5%) from PDAC tissues were assigned to *cHR*, potentially indicative of field cancerization within these specimens.

To validate the strong association between IPMN transcriptomic states and PDAC subtypes, we applied our GSEA and GSVA analyses to the Carpenter *et al.* dataset (**Supplementary Fig. S14A**). The results mirrored our IPMN findings: 1) classical gene upregulation occurred in PanIN relative to NL AOIs, 2) basal-like gene expression was low in NL and PanIN from donor pancreata and high in PDAC, and 3) exocrine genes were highly expressed in donor pancreata and diminished in PDAC. Identical patterns of PDAC molecular subtype expression were seen in the Carpenter dataset (**Supplementary Fig. S14B**). Thus, an independent DSP-RNA dataset precisely recapitulated our observations from IPMN, indicating that the transcriptomic states that characterize IPMN are generalizable across precancerous lesions of the pancreas.

### Trajectory inference predicts bifurcation of early neoplastic lesions into indolent and malignant lineages

Collectively, our gene expression and enrichment analyses exposed greater differences between *cLR* and *cHR* states than between *cLR* and *cNL* or *cHR* and *cNL*. This led us to speculate that IPMN undergoes divergent evolution from an early neoplastic state into indolent (*cLR*) and malignant (*cHR*) lineages rather than linear evolution along a single axis (NL-LGD-HGD-INV). To model this hypothesis *in silico*, we repeated unsupervised clustering to produce a four-cluster model. We identified *cNL*, *cLR*, and *cHR* clusters, and denoted the fourth cluster *cProgenitor* representing the theoretical common predecessor to *cLR* and *cHR* (**Fig. 5A**). Trajectory inference supported the bifurcating lineage model as hypothesized (**Fig. 5B**). We anticipate that this hypothetical model could drive further mechanistic exploration of intrinsic and extrinsic factors that affect lineage fate in early pancreatic neoplasia.

**Figure 5:**
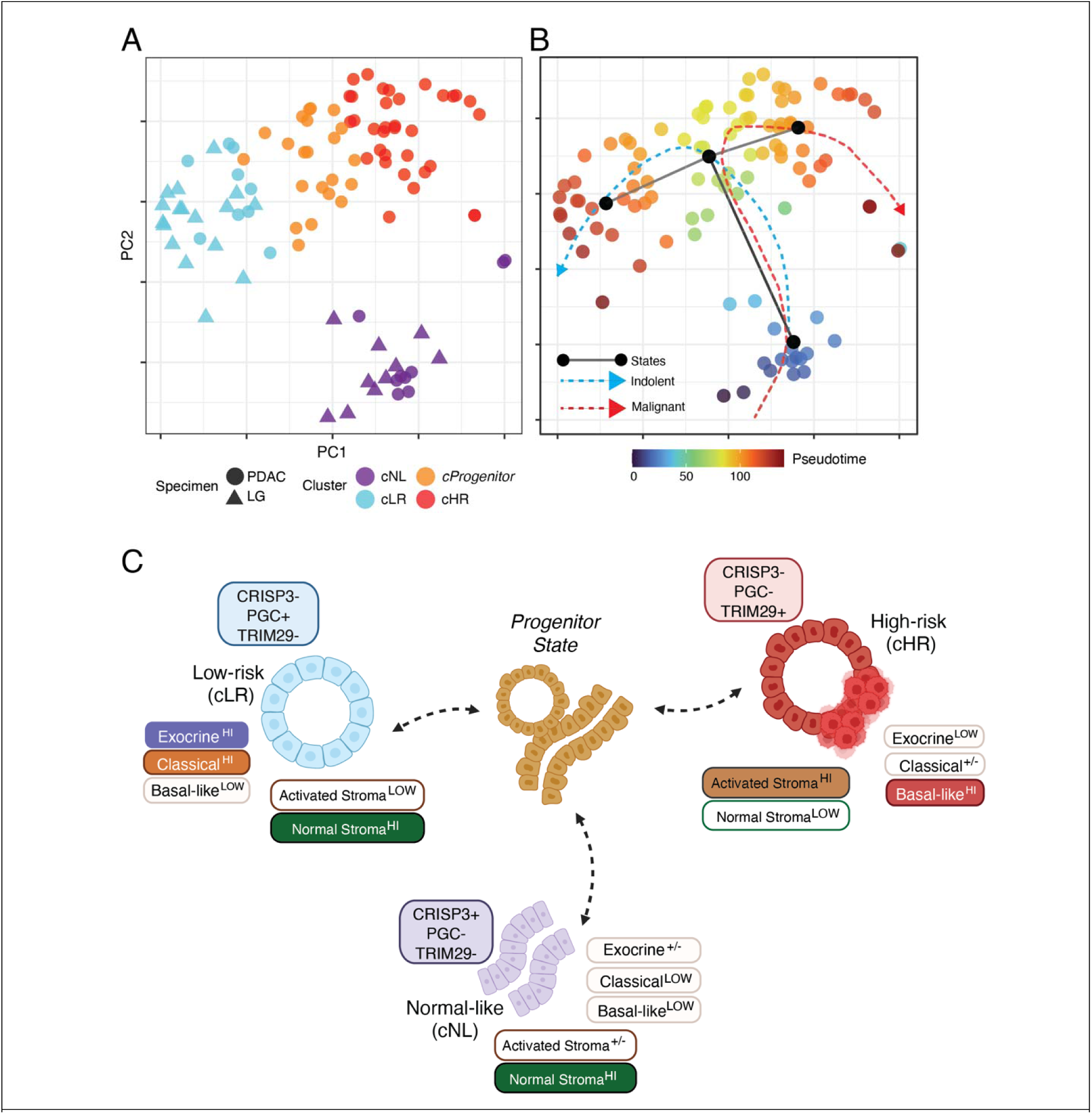
Hypothetical divergent model of IPMN into indolent and malignant lineages. **A)** PCA scatter plot of AOIs showing a four-cluster model of IPMN phenotypes, with the additional cluster identified as *cProgenitor* (orange). **B)** Trajectory inference analysis showing AOI points colored by pseudotime from their computed pseudotime. Black dots interconnected by lines represent transcriptomic state changes. Blue (indolent) and red (malignant) curved lines represent the trajectory of the predicted lineages. **C)** Transcriptomic state model summarizing the results of this study. Normal-like regions (cNL) exhibit prominent expression of ductal cell markers (*CRISP3*), normal stroma, and may or may not express exocrine genes. A hypothetical progenitor or intermediate state between *cLR* and *cHR* characterizes early neoplastic lesions and is associated classical PDAC gene signature upregulation. Low-risk (cLR) lesions exhibit high exocrine and low basal-like expression, whereas high-risk (cHR) lesions exhibit the converse. Gastric neoplasia markers such as *PGC* delineate *cLR*, whereas PDAC markers such as *TRIM29* delineate *cHR*. Stromal expression patterns correlate with the adjacent epithelium. Specifically, *cHR* upregulates and *cLR* downregulates “Activated Stroma” genes relative to *cNL*.

## DISCUSSION

In this study of DSP-RNA profiles across the spectrum of progression from normal ducts to invasive carcinoma arising from IPMN, we uncovered three transcriptomic states identified as “normal-like” (*cNL*), “low-risk” (*cLR*), and “high-risk” (*cHR*) based on their annotated features (**Fig. 5C**). The three states could be characterized by four expression programs: 1) pancreatic ductal cell markers, 2) pancreatic exocrine function, 3) gastric neoplasia, and 4) PDAC-associated genes. Signatures derived from external bulk and single-cell RNA-Seq studies validated and contextualized the IPMN states. We found that the spatial expression profiles of IPMN embodied the exocrine, classical, and basal-like patterns derived from PDAC. An exocrine-basal-like axis explained the differences between *cLR* and *cHR*, and classical PDAC genes distinguished neoplastic (*cLR* and *cHR*) from non-neoplastic (*cNL*) regions. The IFN-γ, TNF-NFκB, and EMT pathways most strongly distinguished *cHR* from *cLR*. Interestingly, *cLR* regions exhibited quiescent inflammatory activity and were more differentiated than their *cNL* counterparts. This was supported by adjacent stromal profiles, where the “activated stroma” signature was lowest in *cLR* and highest in *cHR* AOIs. Evidence of the three states and their characteristics could be recapitulated from the analysis of exclusively benign (NL and LGD) appearing regions, suggesting that malignant potential could be detected without histologic evidence of HGD.

DSP-RNA offered the unique advantage of sampling pathology across a relatively large slide area, allowing us to select and profile the spectrum of histologic grades on a single slide. Alternative spatial transcriptomic solutions may incur batch effects or additional costs to accomplish a similar goal. Moreover, the ability to implicate exocrine expression as integral to the phenotypic characterization of IPMN depended upon precise epithelial cell transcriptome isolation through manual ROI selection and PanCK segmentation, another distinct advantage of DSP-RNA over “spot” based spatial transcriptomics and bulk tissue profiling methods. Deconvolution of imprecise transcriptomic profiles could otherwise lead to misinterpretation of exocrine expression as acinar tissue contamination. We anticipate that the intermixed stromal, immune, and epithelial cell populations of PDAC will continue to challenge transcriptomics analyses performed at greater than single-cell resolution. While DSP-RNA profiles of HGD offered the opportunity to sample the transcriptome of malignant epithelial cells with relatively high purity, the relatively sparse distribution of immune cells within the TME hindered our ability to obtain usable spatial profiles of these areas, and our stromal profiles were frequently convoluted with acinar and islet cell gene expression. Future efforts to profile immune and stromal regions should consider the single-cell spatial platforms now available for studying targeted gene panels.

Our findings add to existing evidence refuting the linear paradigm of carcinogenesis, where epithelial cells advance through sequential stages of dysplasia culminating in malignancy. Instead, we and others suggest that early neoplasia diverges into indolent and malignant lineages (**Fig. 5C**). Supporting this, Burdziak *et al.* observed polarization of pancreatic epithelial cells into indolent and malignant phenotypes in PDAC mouse models by both single-cell RNA-seq and chromatin accessibility studies^12,41^. One explanation for the divergence in transcriptomic states is the possibility that tumors originate from different cell types. Flowers *et al.* and others provided evidence suggesting that mouse acinar cell-derived and ductal cell-derived tumor signatures are enriched in classical and basal-like programs, respectively^42^. Our discovery of an exocrine-basal-like expression axis in humans that distinguishes indolent from malignant regions is consistent with this hypothesis. Tumors originating from ductal cells inherently lack exocrine function, whereas tumors that evolve from acinar-ductal metaplasia may retain some exocrine activity. Further exploration of this hypothesis would require concurrent profiling of pancreatic acini and acinar-ductal metaplasia, which we did not perform in this study.

Alternatively, the transcriptomic phenotypes observed in this study could be associated with specific combinations of genomic mutations. The most mutated genes in IPMN – *KRAS*, *GNAS*, and *RNF43* – are not reliably associated with grade of dysplasia or prognosis^43^. However, mutations in *TP53*, *SMAD4*, *PIK3CA*, and *CDKN2A* were shown to be highly specific, but not sensitive, for HGD/INV IPMN^44,45^. A genomic basis for lineage bifurcation is compatible with the Burdziak *et al.* study, which utilized *Kras*-mutant and *Kras-*mutant/*p53*-mutant mice to model early neoplastic and malignant lineages, respectively. Recently, Braxton *et al*. leveraged CODA, a method for 3D reconstruction of large tissues from serially sectioned H&E images, with regional microdissection to achieved spatial genomic mapping of pancreatic cancer precursors^46,47^. This effort unveiled a preponderance of polyclonal pancreatic intraepithelial neoplasias (PanIN) in tissue slabs from patients with and without associated PDAC. However, correlative spatial transcriptomic profiling of these tissue slabs was not performed. In addition to genomic aberrations, tumor-extrinsic inflammatory injury is associated with lineage plasticity and tumor initiation and must be accounted for in PDAC progression^48^. Hence, methods for combined spatial genomic and transcriptomic profiling are highly anticipated and would be a powerful platform to characterize tumor evolution.

This study reframes the classical and basal-like PDAC signatures as powerful predictors of transcriptomic phenotypes in precancerous lesions. Here, the classical program was activated to varying degrees across all neoplastic lesions, whereas basal-like expression associated with regions classified as “high-risk”. Interpretation of tumor subtyping from this perspective aligns well with prior observations that classical and basal-like states coexist in single cells and across tumors^31,49^. Our observation of basal-like activity as a specific characteristic of malignant lesions motivates the need for therapeutic agents targeting the basal-like program.

The notion of diverging indolent and malignant phenotypes has implications for patient care, which currently depends upon clinical and radiographic risk stratification. Circumstantial evidence supporting the existence of an indolent IPMN subtype continues to amass from observational trials of patients with incidental pancreatic cysts as well as favorable outcomes data from patients who undergo pancreatectomy for benign IPMN^50,51^. This work nominates putative clinical biomarkers that could augment risk stratification. For example, *TRIM29* was upregulated by >4 log2-fold in *cHR* regions, and *PGC* was upregulated by an at least >8 log2- fold in *cLR* regions. Future studies are warranted to assess the clinical potential of these and other markers in pancreatic cyst fluid.

Patients who undergo resection for IPMN have an elevated risk of cancer in the remnant gland, leading to the consideration of IPMN as a field defect that affects the surrounding organ^52^. Indeed, the presence of malignant gene signatures within adjacent NL and LGD regions provides molecular evidence for the concept of “field cancerization” in the pathogenesis of IPMN^53^. Our comparison of NL and LGD regions from patients with and without cancer yielded specific genes implicated in field cancerization that could be leveraged for risk stratification. For example, a pinpoint focus of HGD within a background of “cancerized” benign tissue may be detectable through markers of field cancerization rather than the foci of HGD itself.

In addition to the limitations mentioned above, our study cohort of ten patients may not capture less common transcriptomic states. A crucial clinical conundrum is preoperatively detecting HGD before the development of INV. Importantly, we did not include IPMNs with HGD alone (lacking invasive carcinoma) in this study and cannot assume that these specimens would fall into the high-risk state (*cHR*) defined by our analysis. We also did not include INT specimens, which have distinct transcriptomes and relatively favorable clinical outcomes compared to PB. Future work should leverage this dataset and analysis approach as part of a more comprehensive effort to classify precancerous pancreatic lesions.

In summary, this study examines the histopathology spectrum of IPMN in both low-grade and malignant specimens defines the major transcriptomic subtypes of IPMN. Our transcriptomic characterization, validated through external DSP-RNA datasets and bulk RNA-Seq signatures, supports a model of pancreatic ductal neoplasia in which early progenitors diverge into either indolent or malignant lineages. Notably, we identified a specific gene expression program associated with malignant fate, closely related to basal-like PDAC, which could be detected even in histologically benign (NL or LGD) lesions. We anticipate that our findings will contribute to the development of new molecular biomarkers for risk stratification of IPMN and stimulate further inquiry into the determinants of lineage trajectory.

## Acknowledgments

We thank Antonio Agostini and Carmine Carbone from the Department of Translational Medicine, Catholic University of the Sacred Heart, Rome, Italy for sharing Nanostring GeoMx data and annotations related to GSE229752. We thank Hariharan K. Iyer for thoughtful discussion of the statistical approaches and considerations involved in spatial transcriptomics data normalization. The authors acknowledge funding support from the Marshall and Therese Sonenshine Foundation and NIH R01 CA182076.

## Data and materials availability

Code implementing quality control and normalization can be found on GitHub at https://github.com/mkiyer/stgeomx. Code to reproduce the analysis including all figures and tables generated for this manuscript can be obtained from GitHub at https://github.com/mkiyer/stgeomx_ipmn_wta. Data needed to evaluate the conclusions in the paper are present in the paper and/or the Supplementary Materials.

## Supplementary Figures and Figure Legends

**Supplementary Figure S1:**
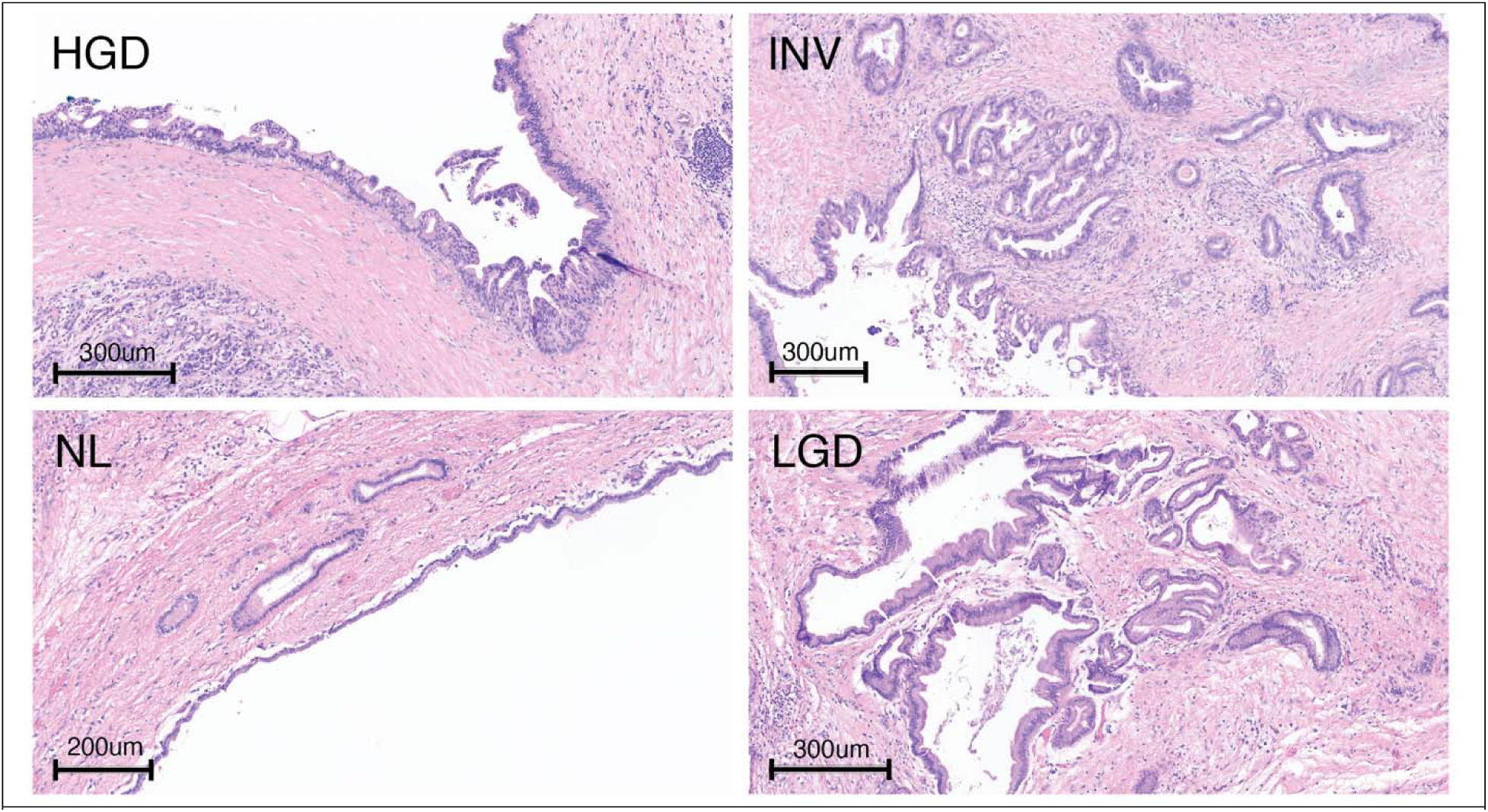
Hematoxylin and eosin (H&E) stained tissue images from patient *p10* corresponding to Figure 1B. Regions of normal duct (NL), low-grade dysplasia (LGD), high-grade dysplasia (HGD), and invasive carcinoma (INV) were selected for digital spatial profiling.

**Supplementary Figure S2:**
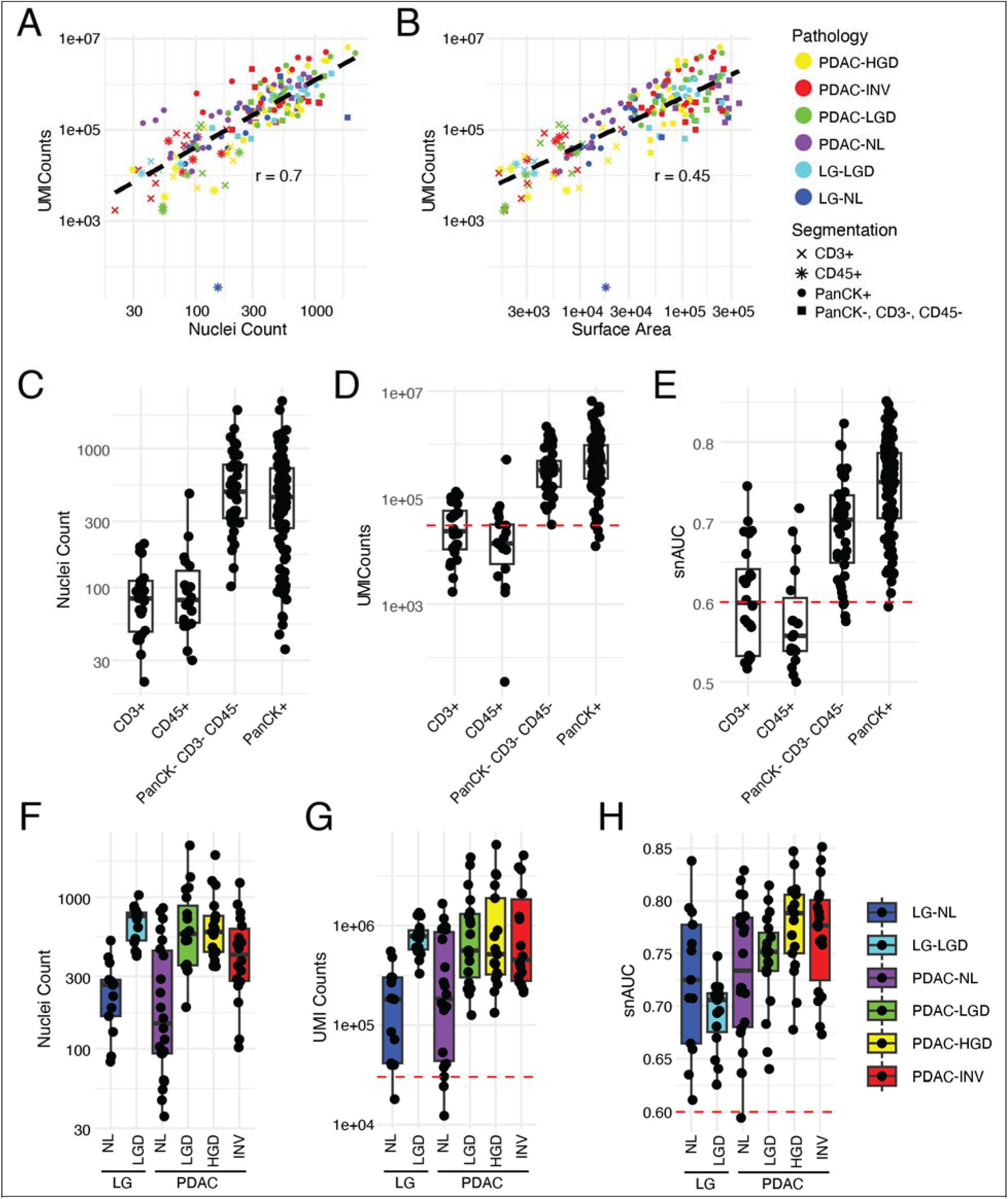
Quality control of DSP-RNA data. **(A,B)** Scatter plots showing correlation between (**A**) AOI nuclei count and UMI counts and (**B**) AOI surface area and UMI counts. **(C, D, E)** Box plots showing relationship between ROI segmentation and **(C)** nuclei count, **(D)** UMI counts, and **(E)** snAUC. **(F, G, H)** Box plots showing relationship between histopathology and **(F)** nuclei count, **(G)** UMI counts, and **(H)** snAUC. Abbreviations: snAUC = signal-to-noise area-under-the-ROC-curve, NL = normal, LGD = low-grade, HGD = high-grade, INV = invasive carcinoma.

**Supplementary Figure S3:**
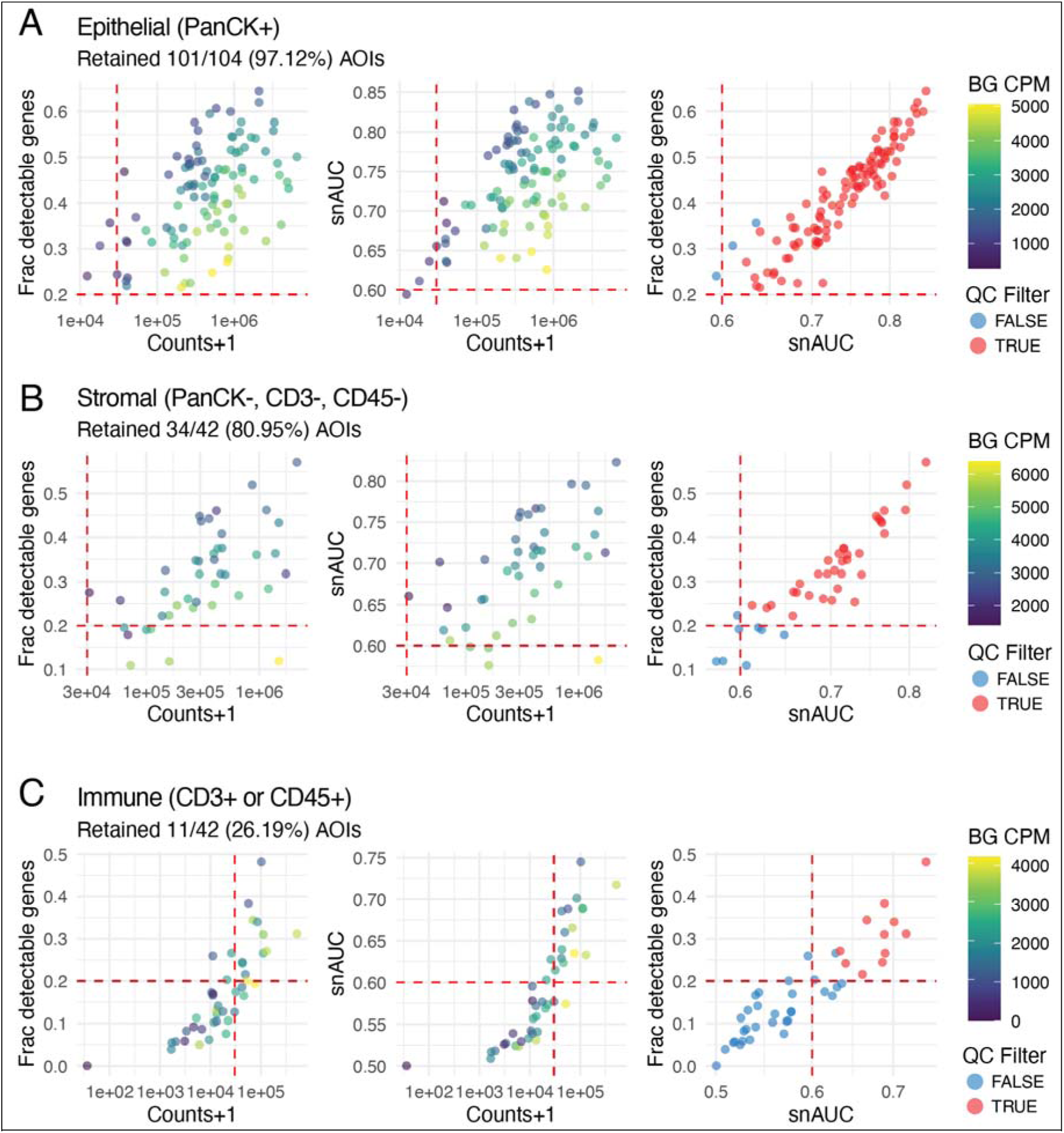
AOI Filtering Criteria. Scatter plots showing (left) counts versus fraction of detectable genes, (middle) counts versus snAUC, and (right) snAUC versus fraction of detectable genes. Dotted red lines indicate filtering cutoffs. **(A)** epithelial (PanCK+) AOIs, **(B)** Stromal (PanCK-, CD3-, CD45-) AOIs, and **(C)** Immune (CD3+ or CD45+) AOIs,

**Supplementary Figure S4:**
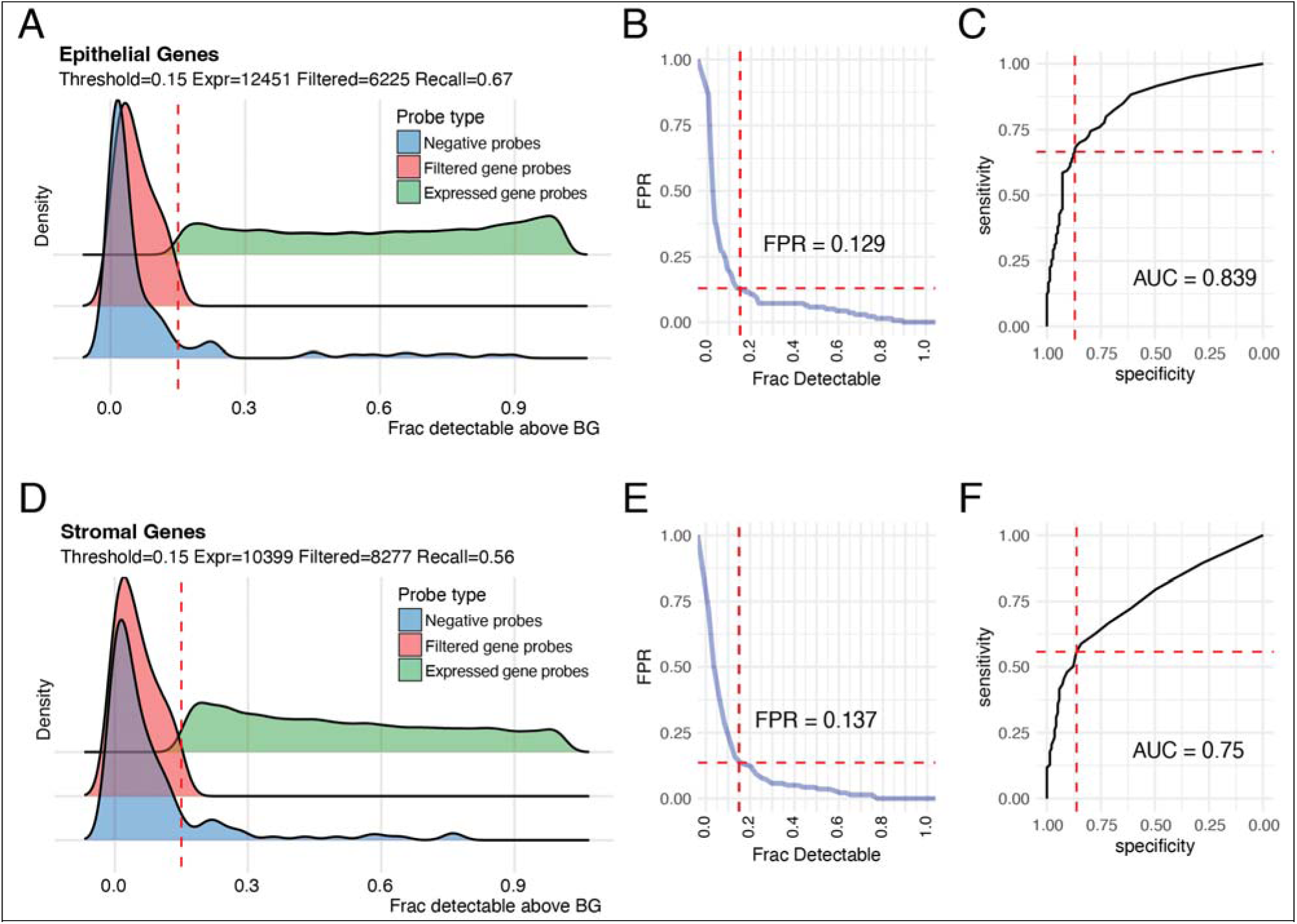
Gene Filtering Criteria. (A-C) Gene filtering for Epithelial AOIs. **(A)** Density plots showing distribution of fraction of detectable probes relative to background probes for each gene. Dotted red line denotes threshold of 0.15 (15% detectable above background) for this dataset, producing expressed and filtered gene probe distributions. **(B)** Plot of false-positive-rate (FPR) for expressed genes relative to fraction of detectable genes, with dotted lines denoting metrics for this dataset. **(C)** Receiver operating characteristic (ROC) curve for detection of expressed versus background probes. **(D-F)** Gene filtering for Stromal AOIs.

**Supplementary Figure S5:**
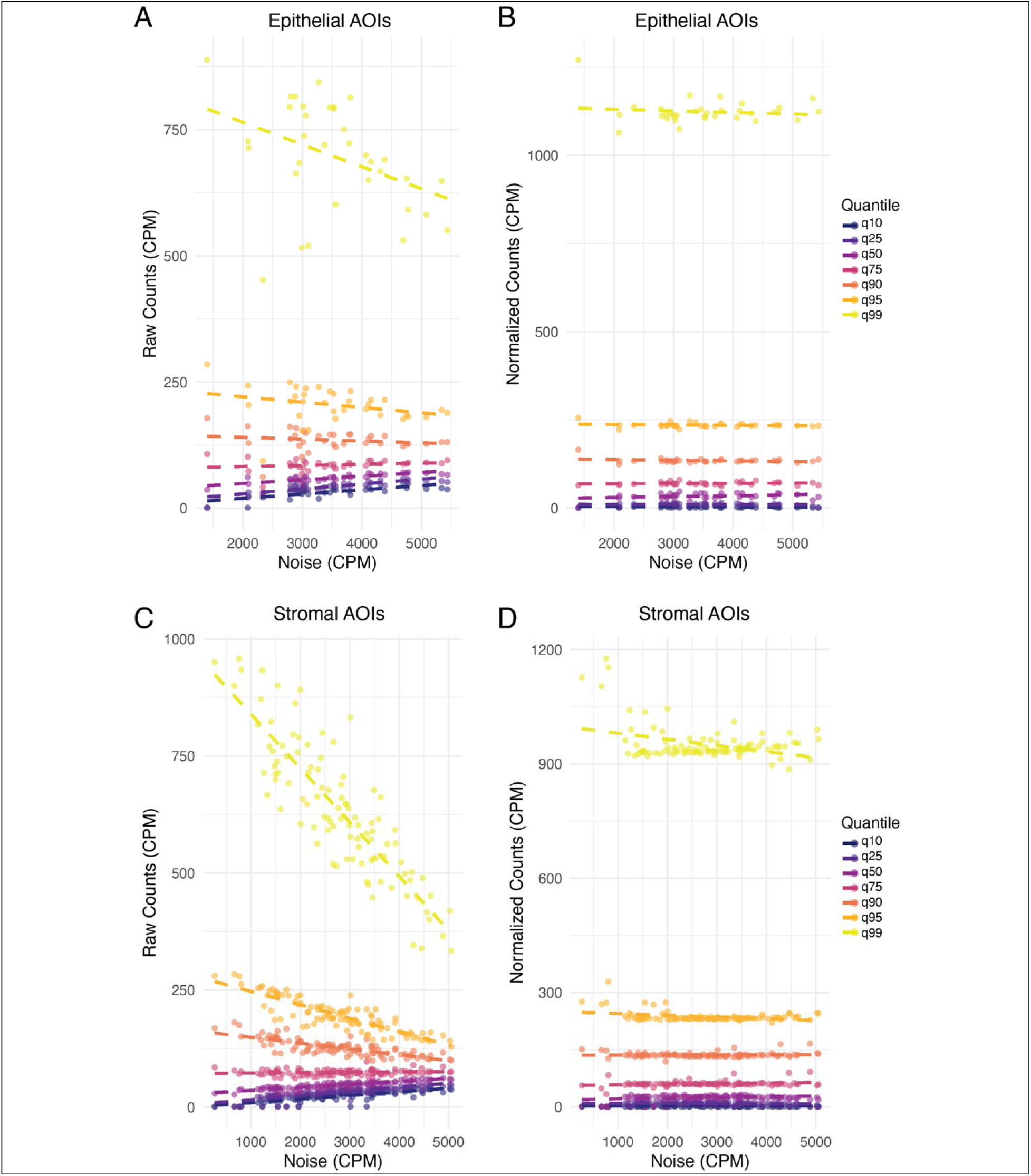
Normalization of DSP-RNA data. Scatter plot of negative probe (background noise) levels (*x* axis) versus gene probes (*y* axis) for each AOI, with quantiles indicated by different colors. **(A)** raw gene probe counts for epithelial AOIs, **(B)** normalized counts-per-million (CPM) for epithelial AOIs, **(C)** raw gene probe counts for stromal AOIs, **(C)** normalized CPM for stromal AOIs

**Supplementary Figure S6:**
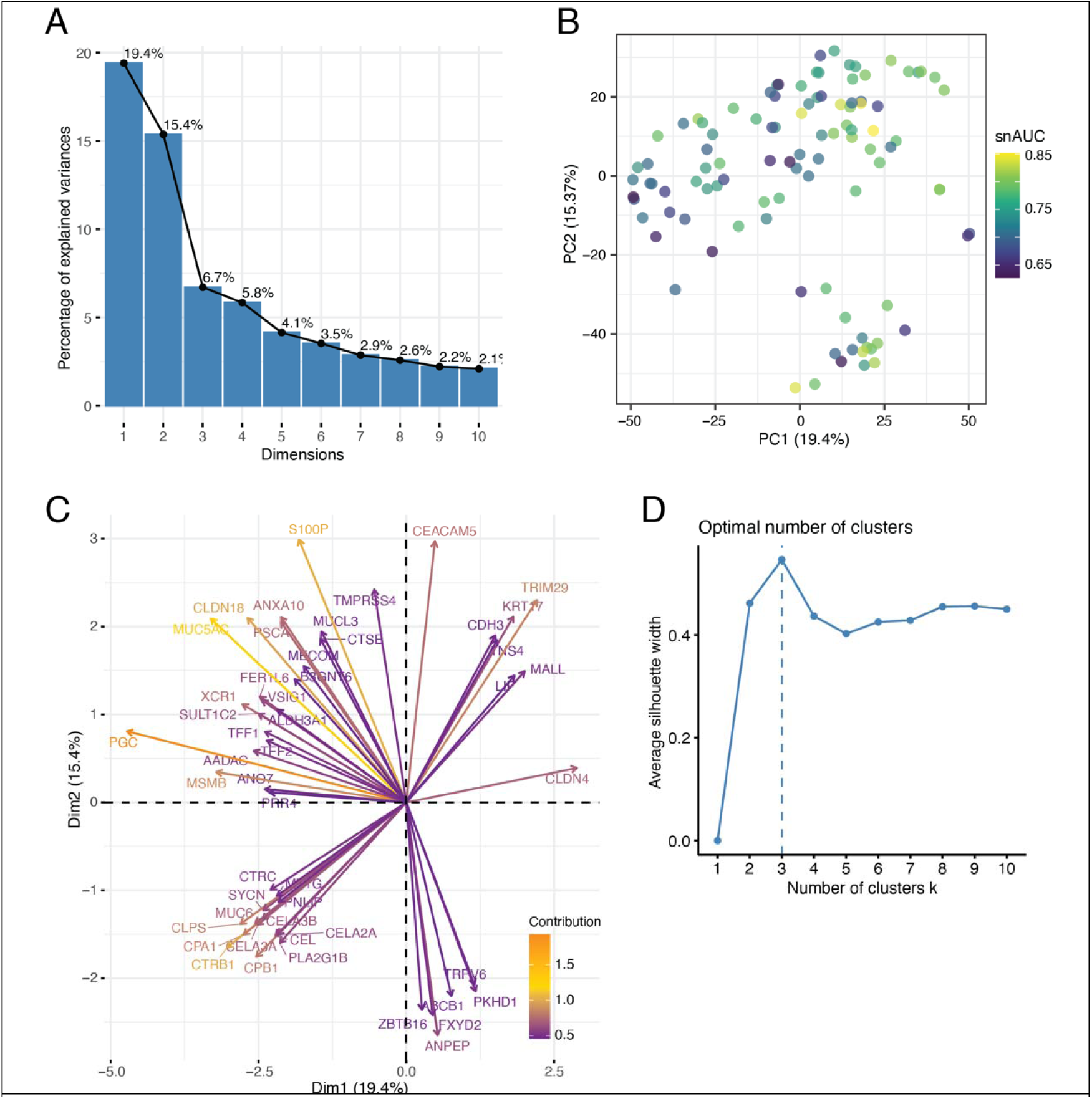
Dimension reduction analysis of DSP-RNA data. **(A)** Bar plot denotes contribution of each principal component (PC) to the percentage of total variance in the dataset. **(B)** PCA feature plot colored by snAUC. **(C)** Graph of genes contributing to the first two PCs. Genes plotted as arrows based on their contribution to PC1 and PC2 and colored by their percentage of explained variance. **(D)** Optimization of unsupervised clustering using the average silhouette width metric. Dotted line shows optimal value.

**Supplementary Figure S7:**
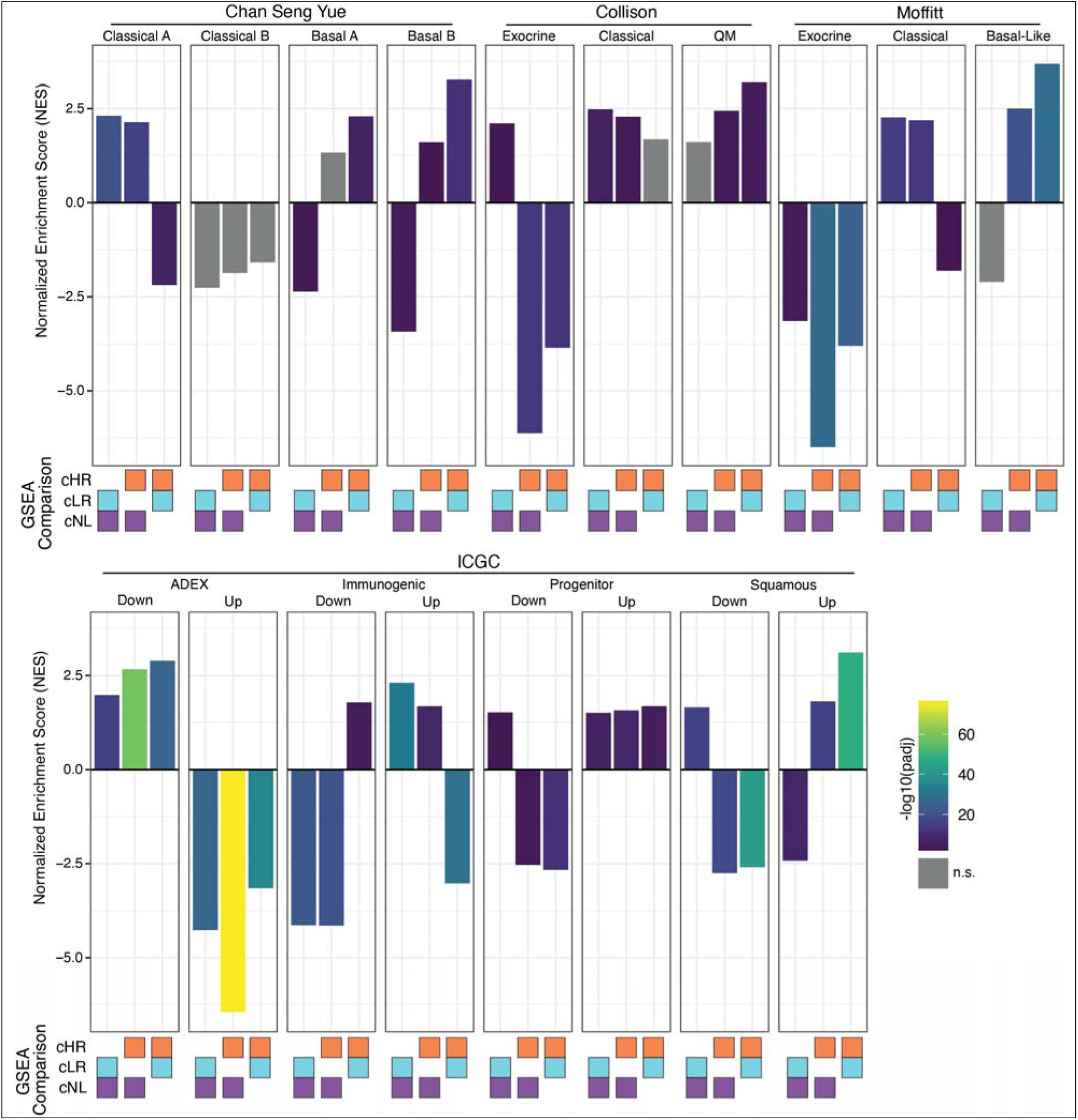
GSEA analysis comparing IPMN clusters to PDAC molecular subtyping studies. Bar height indicates the normalized enrichment score (NES) of each GSEA comparison and bar color denotes the adjusted p-value. Boxes below each bar indicate the ranked list of genes that generated the result (*cLR-vs-cNL*, *cHR- vs-cNL*, or *cHR-vs-cLR*). Sets of plots are grouped by study and gene set.

**Supplementary Figure S8:**
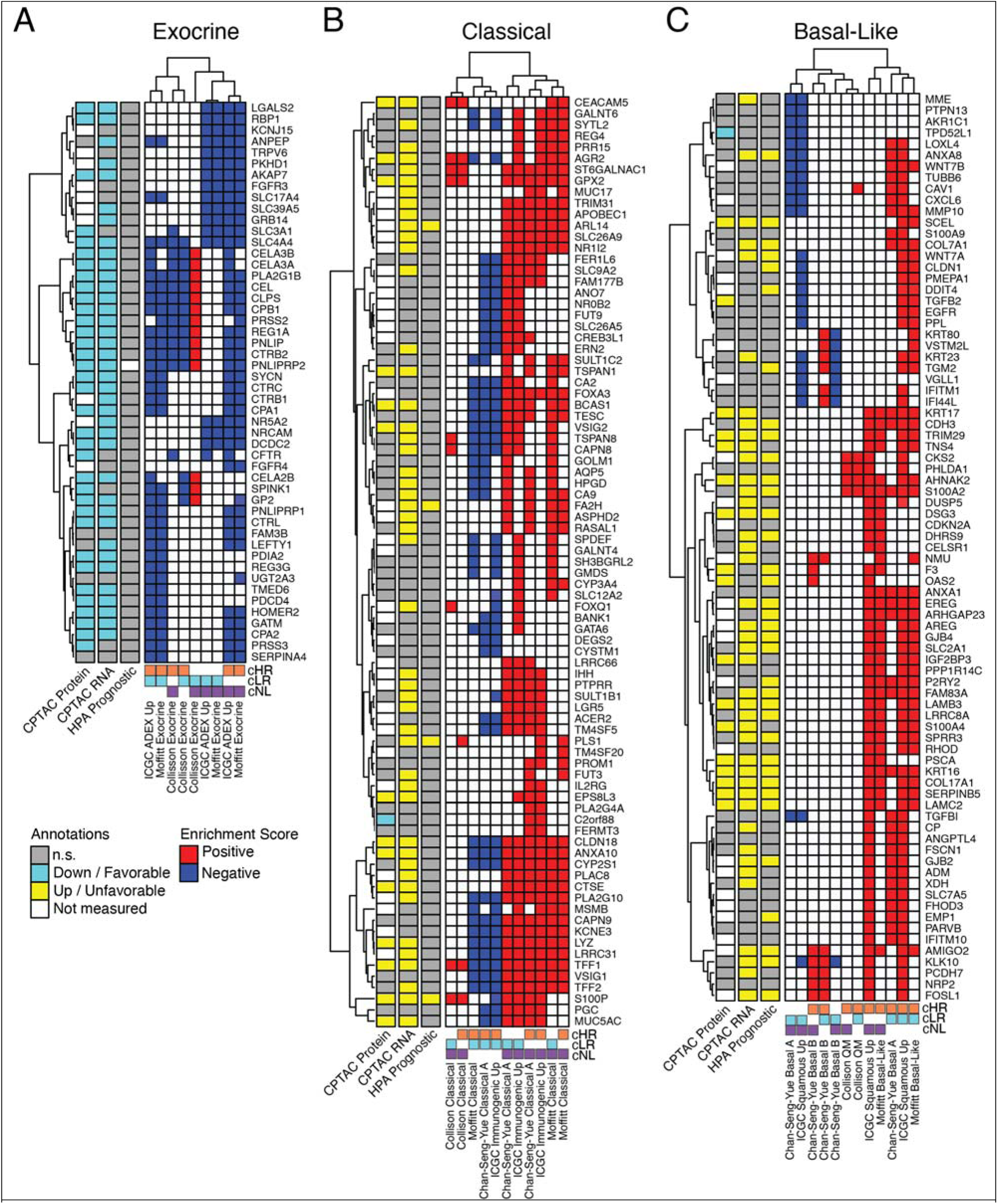
Leading edge analysis of GSEA results. Leading edge heatmap plots for **(A)** exocrine, **(B)** classical, and **(C)** basal-like GSEA analyses. Rows denote enriched genes, and each column represents a GSEA result. Colors indicate positive enrichment (red) or negative enrichment (blue). Row annotations (left) show membership of genes in CPTAC RNA/Protein and HPA Prognostic gene sets for context. Hierarchical clustering is applied to expose common gene subsets.

**Supplementary Figure S9:**
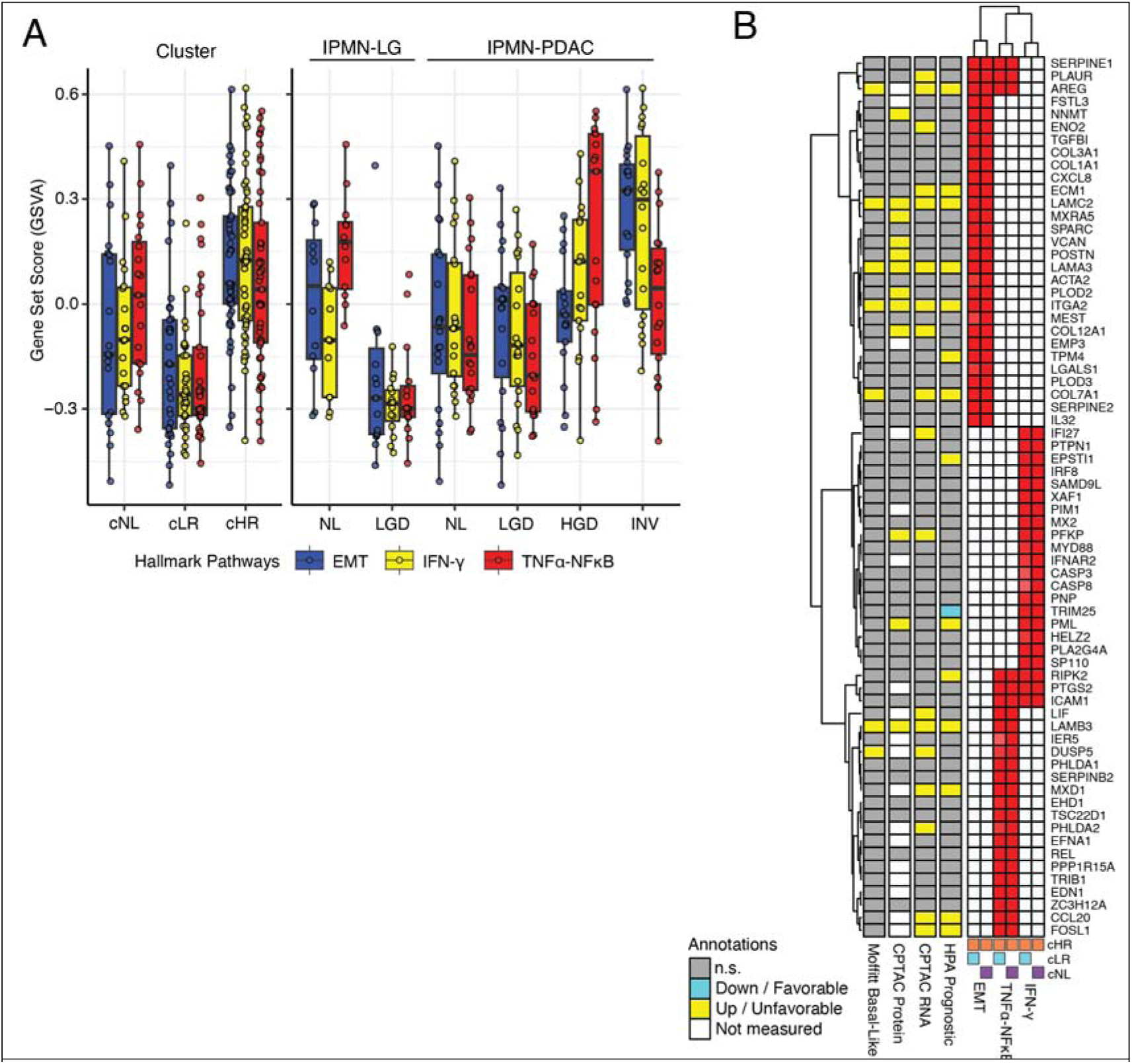
Hallmark pathways associated with IPMN transcriptomic states. **(A)** Boxplots showing the distribution of scores of the top Hallmark pathways for clusters and histopathologic phenotypes. **(B)** Leading edge heatmap plot showing DE genes enriched in EMT, TNF-NFκB, and IFN-γ, pathways for each comparison. Annotations show genes present in CPTAC, HPA, and Moffitt Basal-like gene sets.

**Supplementary Figure S10:**
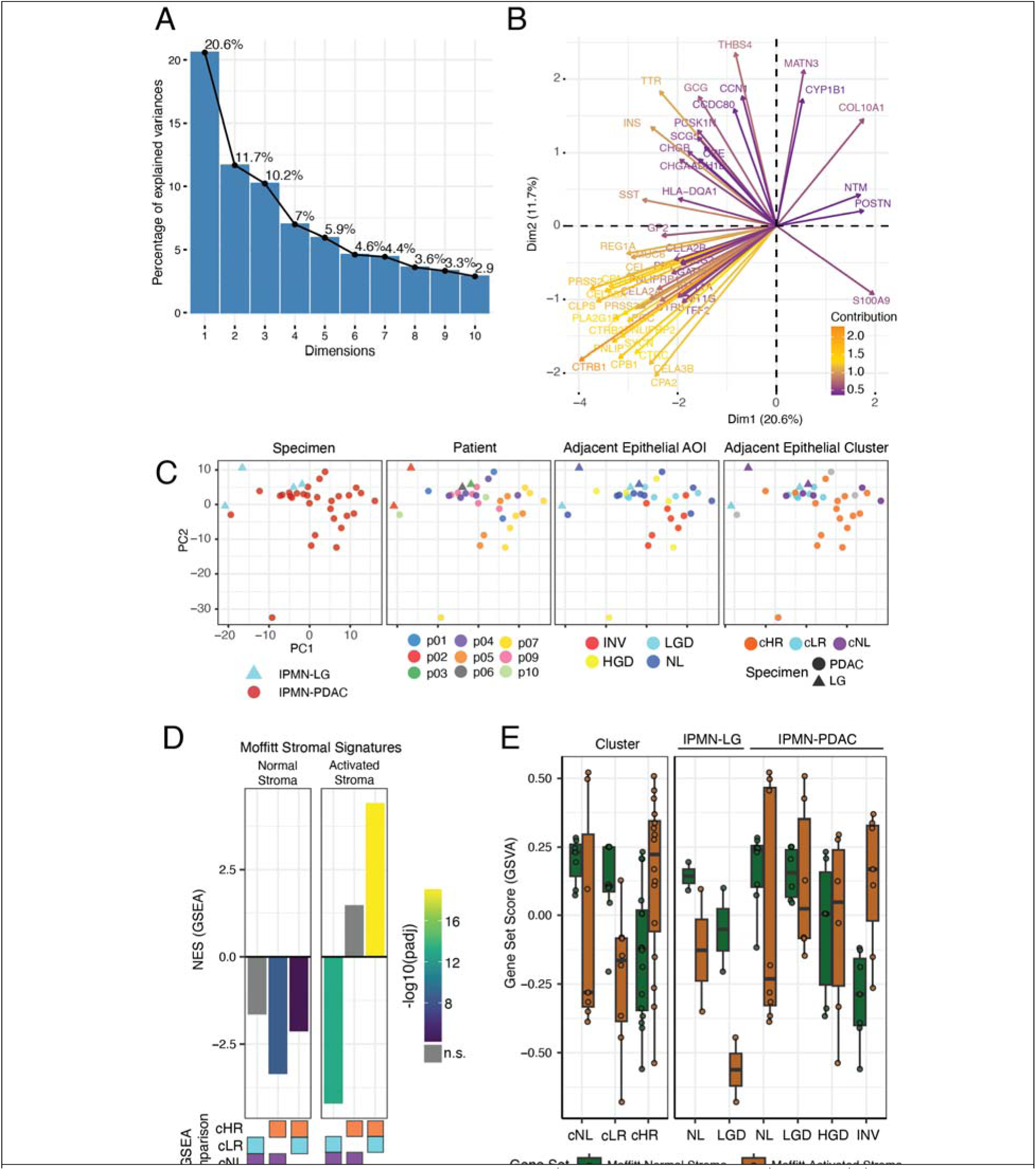
Analysis of adjacent stromal (PanCK-, CD3-, CD45-) AOIs. **(A)** Bar plot of contribution of each principal component (PC) to the percentage of total variance. **(B)** Graph of genes contributing to the first two PCs. Genes plotted as arrows based on their contribution to PC1 and PC2 and colored by their percentage of explained variance. **(C)** Scatter plots of PC1 (*x* axis) versus PC2 (*y* axis) for stromal gene signature colored by specimen, patient, adjacent epithelial histopathology, and adjacent epithelial cluster. **(D)** Bar plots showing results of GSEA analysis of Moffitt stromal signatures, where bar height indicates the normalized enrichment score (NES) of each GSEA comparison and bar color denotes the adjusted p-value. Boxes below each bar indicate the ranked list of genes that generated the result (*cLR-vs-cNL*, *cHR-vs-cNL*, or *cHR-vs-cLR*). **(E)** Boxplots showing the GSVA scores of the Moffitt stromal signatures grouped by cluster and histology.

**Supplementary Figure S11:**
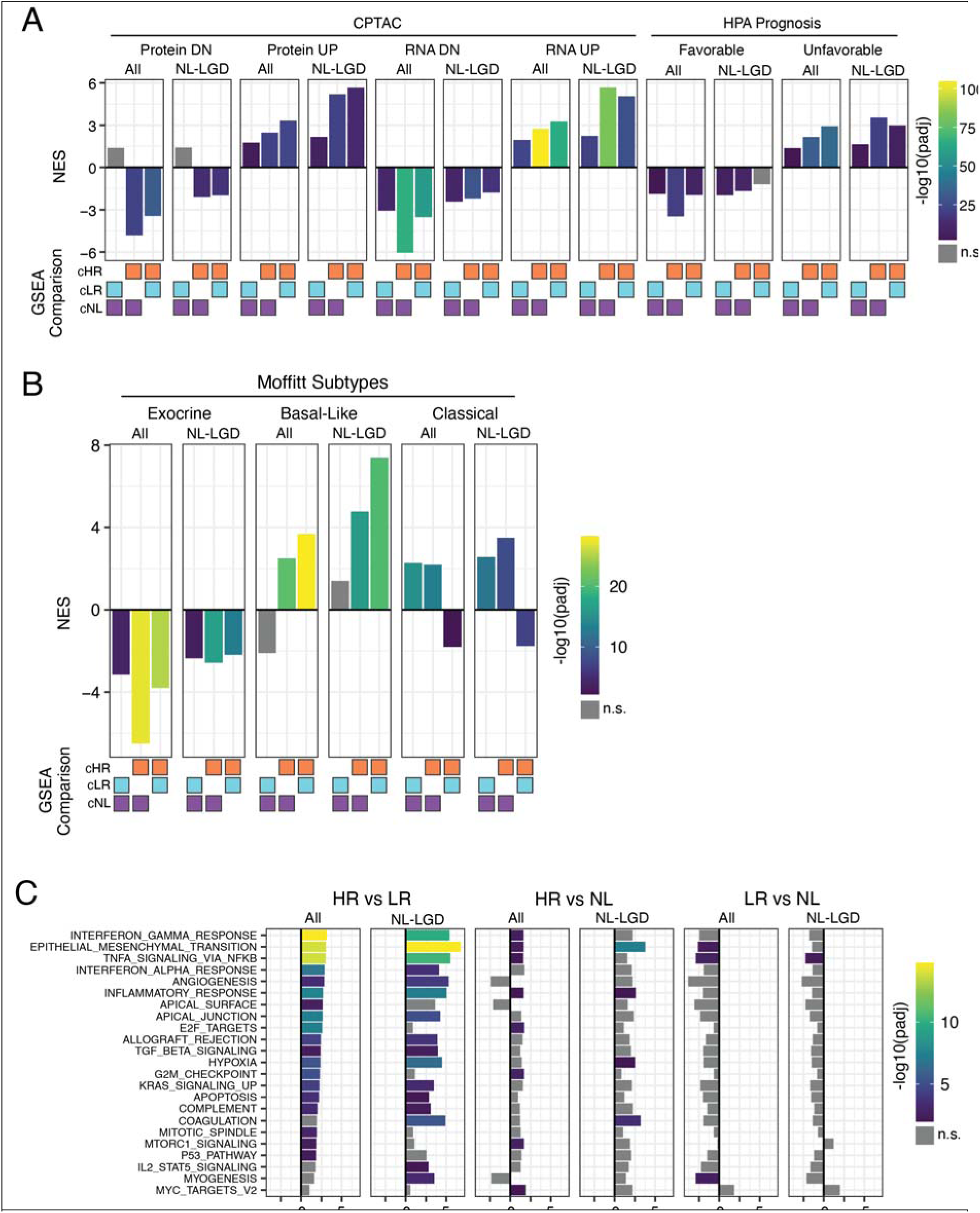
GSEA analyses of NL-LGD AOIs. **(A-C)** Sets of barplots illustrating GSEA results comparing the analysis of *all* AOIs to a separate analysis of just the subset of benign (NL and LGD) ROIs for **(A)** CPTAC and HPA gene sets, **(B)** Moffitt PDAC subtypes, and **(C)** Hallmark molecular pathways. Bar height represents normalized enrichment score, and bar color reflects adjusted p-value. Grey bars are not statistically significant.

**Supplementary Figure S12:**
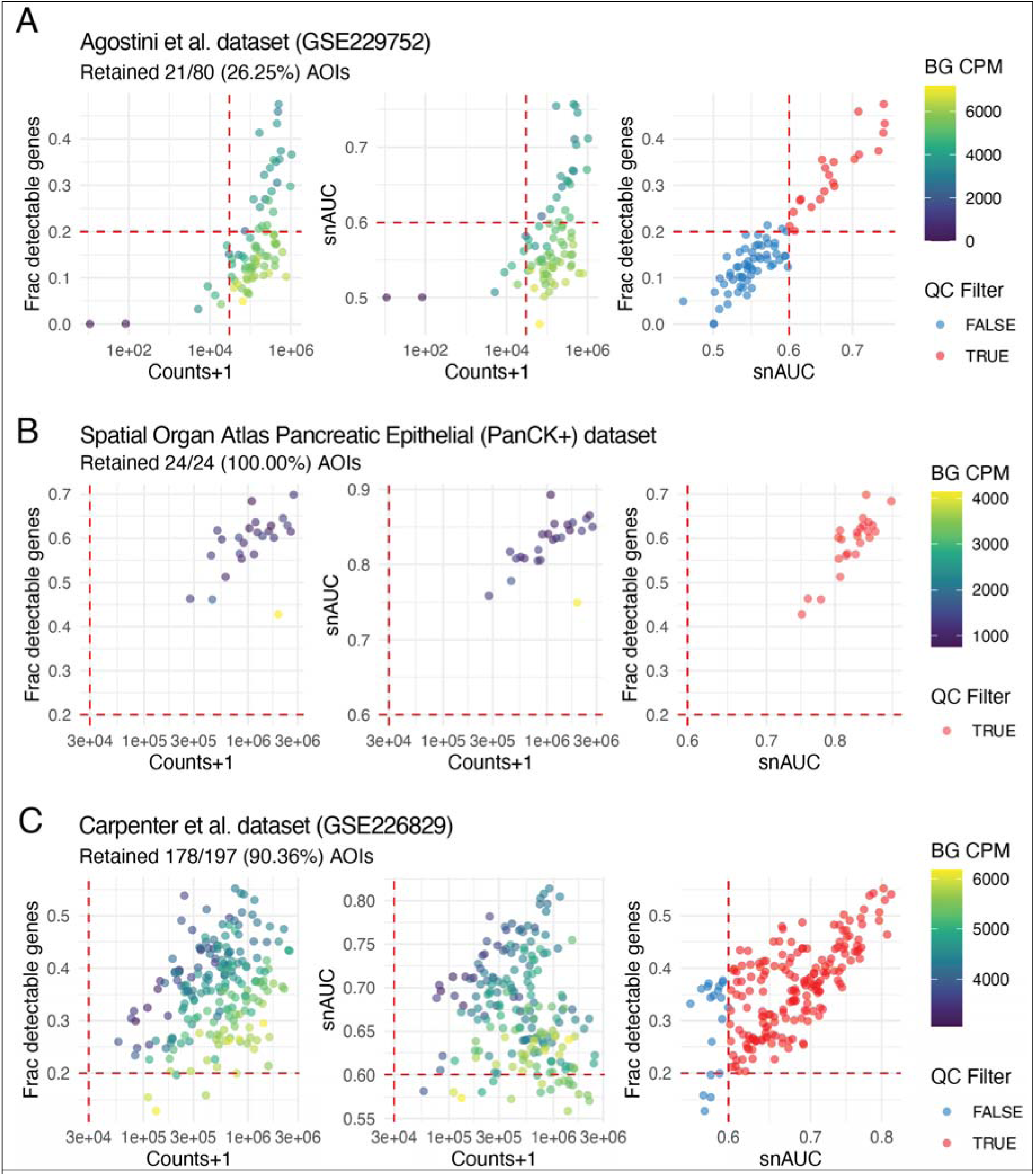
Quality control of external DSP-RNA datasets incorporated for validation. Scatter plots showing (left) counts versus fraction of detectable genes, (middle) counts versus snAUC, and (right) snAUC versus fraction of detectable genes. Dotted red lines indicate filtering cutoffs. **(A)** Agostini *et al*. (GSE229752) **(B)** Spatial Organ Atlas **(C)** Carpenter *et al.* (GSE226829).

**Supplementary Figure S13:**
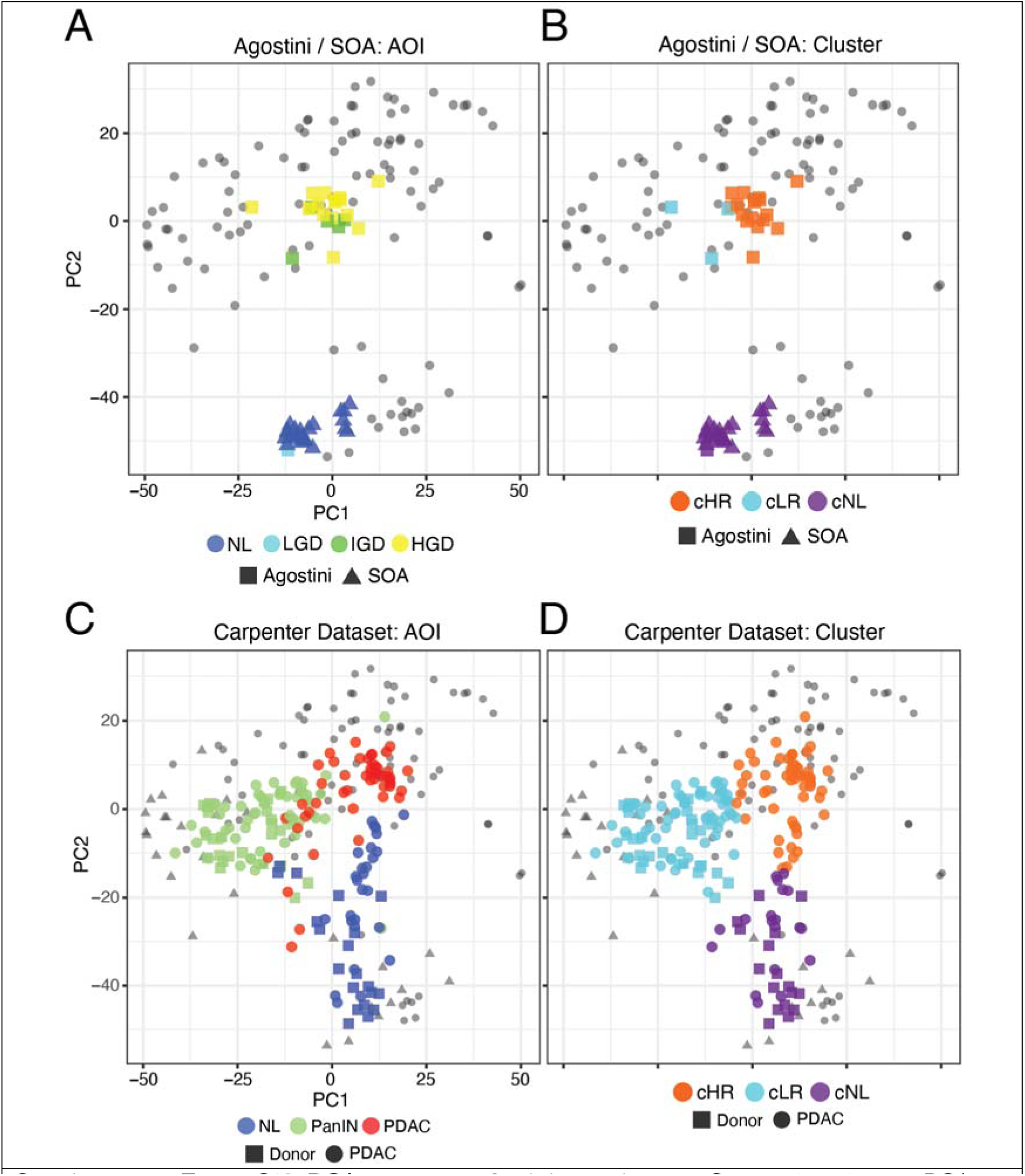
PCA projection of validation datasets. Scatter plots showing PCA projections of the external validation datasets (test data) onto the PCs defined by the IPMN dataset (training data). **(A, B)** Agostini *et al.* and Spatial Organ Atlas (SOA) datasets. Points colored by **(A)** histopathology, **(B)** imputed cluster. **(C, D**) Carpenter *et al.* dataset. Points colored by **(C)** histopathology and **(D)** imputed cluster assignment. Abbreviations: SOA = Spatial Organ Atlas, Agostini = Agostini *et al.* tissue microarray dataset, NL = normal ductal epithelium, LGD = lowgrade dysplasia, IGD = intermediate-grade/borderline dysplasia, HGD = high-grade dysplasia, *cHR* = “high-risk” cluster, *cLR* = “low-risk” cluster, *cNL* = “normal-like” cluster.

**Supplementary Figure S14:**
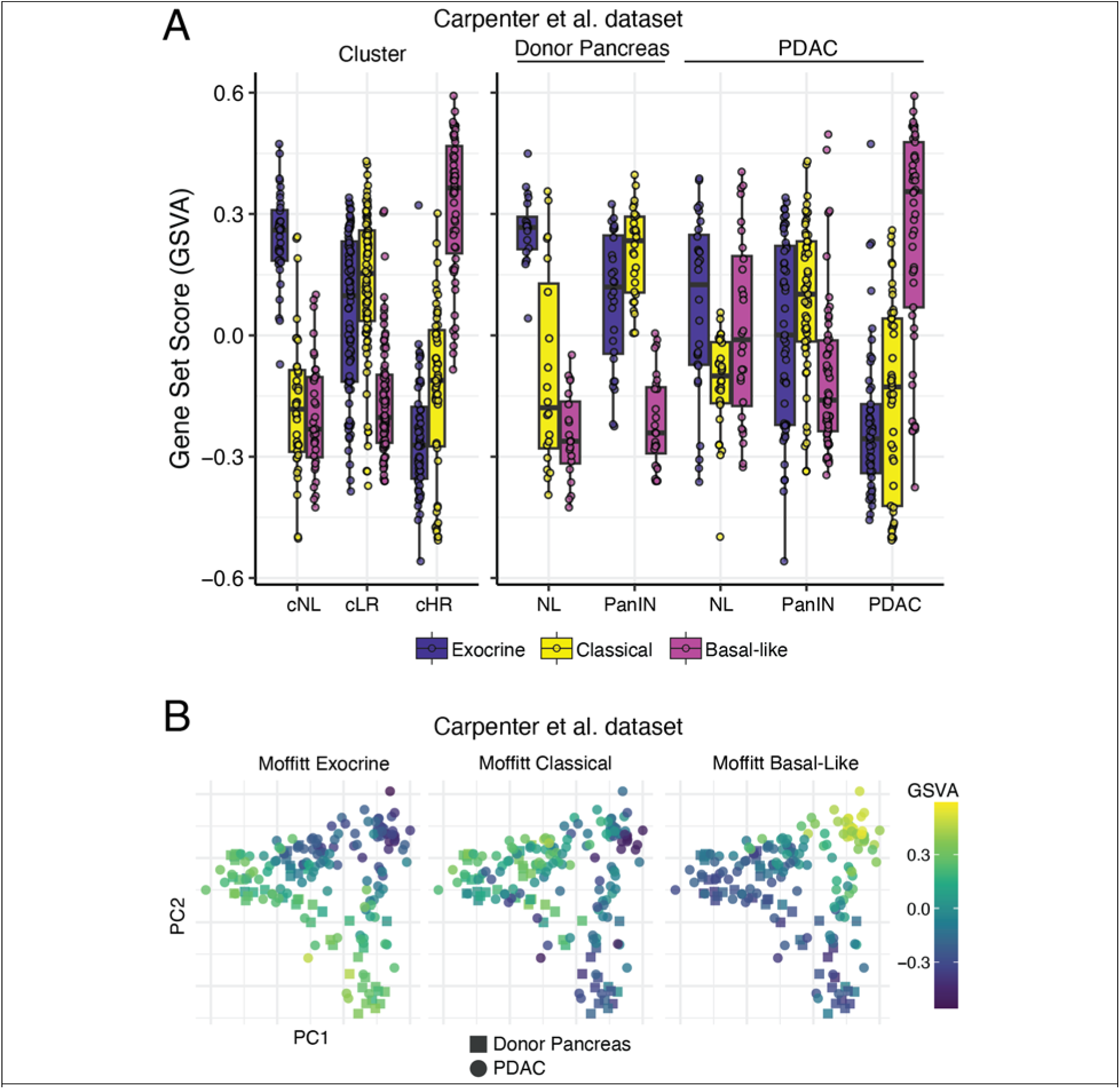
GSEA and GSVA of the Carpenter *et al.* dataset. **(A)** Boxplots showing Moffitt exocrine, classical, and basal-like GSVA scores for AOIs grouped by cluster and histopathology. **(B)** PCA scatter plots colored by the GSVA score of the Moffitt exocrine, classical and basal-like subtypes.

## Notes

### Competing Interest Statement

The authors have declared no competing interest.

## References

1. Siegel RL, Giaquinto AN, Jemal A. Cancer statistics, 2024. CA Cancer J Clin. 2024;74(1):12-49. doi:10.3322/caac.21820

2. Tanaka M, Fernández-del Castillo C, Kamisawa T, et al. Revisions of international consensus Fukuoka guidelines for the management of IPMN of the pancreas. Pancreatology. 2017;17(5):738–753. doi:10.1016/j.pan.2017.07.007

3. Srinivasan N, Teo JY, Chin YK, et al. Systematic review of the clinical utility and validity of the Sendai and Fukuoka Consensus Guidelines for the management of intraductal papillary mucinous neoplasms of the pancreas. HPB. 2018;20(6):497–504. doi:10.1016/j.hpb.2018.01.009

4. Lucocq J, Hawkyard J, Robertson FP, et al. Risk of Recurrence after Surgical Resection for Adenocarcinoma Arising from Intraductal Papillary Mucinous Neoplasia (IPMN) with Patterns of Distribution and Treatment: An International, Multicentre, Observational Study. Ann Surg.:10.1097/SLA.0000000000006144. doi:10.1097/SLA.0000000000006144

5. Winter JM, Jiang W, Basturk O, et al. Recurrence and survival following resection of small IPMN-associated carcinomas (≤ 20 mm invasive component): A multi-institutional analysis. Ann Surg. 2016;263(4):793–801. doi:10.1097/SLA.0000000000001319

6. Sato N, Fukushima N, Maitra A, et al. Gene Expression Profiling Identifies Genes Associated with Invasive Intraductal Papillary Mucinous Neoplasms of the Pancreas. Am J Pathol. 2004;164(3):903–914.

7. Bernard V, Semaan A, Huang J, et al. Single-Cell Transcriptomics of Pancreatic Cancer Precursors Demonstrates Epithelial and Microenvironmental Heterogeneity as an Early Event in Neoplastic Progression. Clin Cancer Res. 2019;25(7):2194–2205. doi:10.1158/1078-0432.CCR-18-1955

8. Wood LD, Adsay NV, Basturk O, et al. Systematic review of challenging issues in pathology of intraductal papillary mucinous neoplasms. Pancreatology. 2023;23(7):878–891. doi:10.1016/j.pan.2023.08.002

9. Iyer MK, Shi C, Eckhoff AM, Fletcher A, Nussbaum DP, Allen PJ. Digital spatial profiling of intraductal papillary mucinous neoplasms: Toward a molecular framework for risk stratification. Sci Adv. 9(11):eade4582. doi:10.1126/sciadv.ade4582

10. Sans M, Makino Y, Min J, et al. Spatial Transcriptomics of Intraductal Papillary Mucinous Neoplasms of the Pancreas Identifies NKX6-2 as a Driver of Gastric Differentiation and Indolent Biological Potential. Cancer Discov. 2023;13(8):1844–1861. doi:10.1158/2159-8290.CD-22-1200

11. Agostini A, Piro G, Inzani F, et al. Identification of spatially-resolved markers of malignant transformation in Intraductal Papillary Mucinous Neoplasms. Nat Commun. 2024;15(1):2764. doi:10.1038/s41467-024-46994-2

12. Burdziak C, Alonso-Curbelo D, Walle T, et al. Epigenetic plasticity cooperates with cell-cell interactions to direct pancreatic tumorigenesis. Science. 2023;380(6645):eadd5327. doi:10.1126/science.add5327

13. Robin X, Turck N, Hainard A, et al. pROC: an open-source package for R and S+ to analyze and compare ROC curves. BMC Bioinformatics. 2011;12(1):77. doi:10.1186/1471-2105-12-77

14. Bolstad BM, Irizarry RA, Åstrand M, Speed TP. A comparison of normalization methods for high density oligonucleotide array data based on variance and bias. Bioinformatics. 2003;19(2):185–193. doi:10.1093/bioinformatics/19.2.185

15. Carpenter ES, Elhossiny AM, Kadiyala P, et al. Analysis of Donor Pancreata Defines the Transcriptomic Signature and Microenvironment of Early Neoplastic Lesions. Cancer Discov. 2023;13(6):1324–1345. doi:10.1158/2159-8290.CD-23-0013

16. Gentleman RC, Carey VJ, Bates DM, et al. Bioconductor: open software development for computational biology and bioinformatics. Genome Biol. 2004;5(10):R80. doi:10.1186/gb-2004-5-10-r80

17. Ritchie ME, Phipson B, Wu D, et al. limma powers differential expression analyses for RNA-sequencing and microarray studies. Nucleic Acids Res. 2015;43(7):e47. doi:10.1093/nar/gkv007

18. Cao L, Huang C, Zhou DC, et al. Proteogenomic Characterization of Pancreatic Ductal Adenocarcinoma. Cell. 2021;184(19):5031–5052.e26. doi:10.1016/j.cell.2021.08.023

19. Uhlen M, Zhang C, Lee S, et al. A pathology atlas of the human cancer transcriptome. Science. 2017;357(6352):eaan2507. doi:10.1126/science.aan2507

20. Torre-Healy LA, Kawalerski RR, Oh K, et al. Open-source curation of a pancreatic ductal adenocarcinoma gene expression analysis platform (pdacR) supports a two-subtype model. Commun Biol. 2023;6(1):1–10. doi:10.1038/s42003-023-04461-6

21. Liberzon A, Birger C, Thorvaldsdóttir H, Ghandi M, Mesirov JP, Tamayo P. The Molecular Signatures Database (MSigDB) hallmark gene set collection. Cell Syst. 2015;1(6):417–425. doi:10.1016/j.cels.2015.12.004

22. Korotkevich G, Sukhov V, Budin N, Shpak B, Artyomov MN, Sergushichev A. Fast gene set enrichment analysis. Published online February 1, 2021:060012. doi:10.1101/060012

23. Hänzelmann S, Castelo R, Guinney J. GSVA: gene set variation analysis for microarray and RNA-Seq data. BMC Bioinformatics. 2013;14(1):7. doi:10.1186/1471-2105-14-7

24. Street K, Risso D, Fletcher RB, et al. Slingshot: cell lineage and pseudotime inference for single-cell transcriptomics. BMC Genomics. 2018;19(1):477. doi:10.1186/s12864-018-4772-0

25. Yonezawa S, Higashi M, Yamada N, Yokoyama S, Goto M. Significance of mucin expression in pancreatobiliary neoplasms. J Hepato-Biliary-Pancreat Sci. 2010;17(2):108–124. doi:10.1007/s00534-009-0174-7

26. Sun H, Dai X, Han B. TRIM29 as a Novel Biomarker in Pancreatic Adenocarcinoma. Dis Markers. 2014;2014(1):317817. doi:10.1155/2014/317817

27. Carpenter ES, Elhossiny AM, Kadiyala P, et al. Analysis of Donor Pancreata Defines the Transcriptomic Signature and Microenvironment of Early Neoplastic Lesions. Cancer Discov. 2023;13(6):1324–1345. doi:10.1158/2159-8290.CD-23-0013

28. Moffitt RA, Marayati R, Flate EL, et al. Virtual microdissection identifies distinct tumor- and stroma-specific subtypes of pancreatic ductal adenocarcinoma. Nat Genet. 2015;47(10):1168–1178. doi:10.1038/ng.3398

29. Collisson EA, Sadanandam A, Olson P, et al. Subtypes of pancreatic ductal adenocarcinoma and their differing responses to therapy. Nat Med. 2011;17(4):500–503. doi:10.1038/nm.2344

30. Bailey P, Chang DK, Nones K, et al. Genomic analyses identify molecular subtypes of pancreatic cancer. Nature. 2016;531(7592):47-52. doi:10.1038/nature16965

31. Chan-Seng-Yue M, Kim JC, Wilson GW, et al. Transcription phenotypes of pancreatic cancer are driven by genomic events during tumor evolution. Nat Genet. 2020;52(2):231–240. doi:10.1038/s41588-019-0566-9

32. Rashid NU, Peng XL, Jin C, et al. Purity Independent Subtyping of Tumors (PurIST), A Clinically Robust, Single-sample Classifier for Tumor Subtyping in Pancreatic Cancer. Clin Cancer Res. 2020;26(1):82–92. doi:10.1158/1078-0432.CCR-19-1467

33. Subramanian A, Tamayo P, Mootha VK, et al. Gene set enrichment analysis: A knowledge-based approach for interpreting genome-wide expression profiles. Proc Natl Acad Sci. 2005;102(43):15545–15550. doi:10.1073/pnas.0506580102

34. O’Kane GM, Grünwald BT, Jang GH, et al. GATA6 Expression Distinguishes Classical and Basal-like Subtypes in Advanced Pancreatic Cancer. Clin Cancer Res. 2020;26(18):4901–4910. doi:10.1158/1078-0432.CCR-19-3724

35. Shockley KE, To B, Chen W, Lozanski G, Cruz-Monserrate Z, Krishna SG. The Role of Genetic, Metabolic, Inflammatory, and Immunologic Mediators in the Progression of Intraductal Papillary Mucinous Neoplasms to Pancreatic Adenocarcinoma. Cancers. 2023;15(6):1722. doi:10.3390/cancers15061722

36. Busser B, Sancey L, Brambilla E, Coll JL, Hurbin A. The multiple roles of amphiregulin in human cancer. Biochim Biophys Acta BBA - Rev Cancer. 2011;1816(2):119–131. doi:10.1016/j.bbcan.2011.05.003

37. Erice O, Narayanan S, Feliu I, et al. LAMC2 Regulates Key Transcriptional and Targetable Effectors to Support Pancreatic Cancer Growth. Clin Cancer Res. 2023;29(6):1137–1154. doi:10.1158/1078-0432.CCR-22-0794

38. Chan A, Prassas I, Dimitromanolakis A, et al. Validation of Biomarkers That Complement CA19.9 in Detecting Early Pancreatic Cancer. Clin Cancer Res. 2014;20(22):5787–5795. doi:10.1158/1078-0432.CCR-14-0289

39. Curtius K, Wright NA, Graham TA. An evolutionary perspective on field cancerization. Nat Rev Cancer. 2018;18(1):19–32. doi:10.1038/nrc.2017.102

40. Basturk O, Hong SM, Wood LD, et al. A REVISED CLASSIFICATION SYSTEM AND RECOMMENDATIONS FROM THE BALTIMORE CONSENSUS MEETING FOR NEOPLASTIC PRECURSOR LESIONS IN THE PANCREAS. Am J Surg Pathol. 2015;39(12):1730–1741. doi:10.1097/PAS.0000000000000533

41. Alonso-Curbelo D, Ho YJ, Burdziak C, et al. A gene–environment-induced epigenetic program initiates tumorigenesis. Nature. 2021;590(7847):642-648. doi:10.1038/s41586-020-03147-x

42. Flowers BM, Xu H, Mulligan AS, et al. Cell of Origin Influences Pancreatic Cancer Subtype. Cancer Discov. 2021;11(3):660–677. doi:10.1158/2159-8290.CD-20-0633

43. Tan MC, Basturk O, Brannon AR, et al. GNAS and KRAS Mutations Define Separate Progression Pathways in Intraductal Papillary Mucinous Neoplasm-Associated Carcinoma. J Am Coll Surg. 2015;220(5):845–854.e1. doi:10.1016/j.jamcollsurg.2014.11.029

44. Pflüger MJ, Jamouss KT, Afghani E, et al. Predictive ability of pancreatic cyst fluid biomarkers: A systematic review and meta-analysis. Pancreatology. 2023;23(7):868–877. doi:10.1016/j.pan.2023.05.005

45. Ohtsuka T, Fernandez-del Castillo C, Furukawa T, et al. International evidence-based Kyoto guidelines for the management of intraductal papillary mucinous neoplasm of the pancreas. Pancreatology. 2024;24(2):255–270. doi:10.1016/j.pan.2023.12.009

46. Kiemen AL, Braxton AM, Grahn MP, et al. CODA: quantitative 3D reconstruction of large tissues at cellular resolution. Nat Methods. 2022;19(11):1490–1499. doi:10.1038/s41592-022-01650-9

47. Braxton AM, Kiemen AL, Grahn MP, et al. 3D genomic mapping reveals multifocality of human pancreatic precancers. Nature. 2024;629(8012):679-687. doi:10.1038/s41586-024-07359-3

48. Falvo DJ, Grimont A, Zumbo P, et al. A reversible epigenetic memory of inflammatory injury controls lineage plasticity and tumor initiation in the mouse pancreas. Dev Cell. 2023;58(24):2959–2973.e7. doi:10.1016/j.devcel.2023.11.008

49. Raghavan S, Winter PS, Navia AW, et al. Microenvironment drives cell state, plasticity, and drug response in pancreatic cancer. Cell. 2021;184(25):6119–6137.e26. doi:10.1016/j.cell.2021.11.017

50. Vaalavuo Y, Vornanen M, Ahola R, et al. Long-term (10-year) outcomes and prognostic factors in resected intraductal papillary mucinous neoplasm tumors in Finland: A nationwide retrospective study. Surgery. 2023;174(1):75–82. doi:10.1016/j.surg.2023.02.006

51. de la Fuente J, Chatterjee A, Lui J, et al. Long-Term Outcomes and Risk of Pancreatic Cancer in Intraductal Papillary Mucinous Neoplasms. JAMA Netw Open. 2023;6(10):e2337799. doi:10.1001/jamanetworkopen.2023.37799

52. Pollini T, Wong P, Maker AV. The Landmark Series: Intraductal Papillary Mucinous Neoplasms of the Pancreas—From Prevalence to Early Cancer Detection. Ann Surg Oncol. 2023;30(3):1453–1462. doi:10.1245/s10434-022-12870-w

53. Curtius K, Wright NA, Graham TA. An evolutionary perspective on field cancerization. Nat Rev Cancer. 2018;18(1):19–32. doi:10.1038/nrc.2017.102

